# The gut microbiota-derived metabolite indole regulates cytoskeletal functions and virulence in *Entamoeba histolytica*

**DOI:** 10.1101/2025.08.01.668081

**Authors:** Eva Zanditenas, Meirav Trebicz Geffen, Yumiko Saito-Nakano, Seiki Kobayashi, Hajime Hisaeda, Tomoyoshi Nozaki, Yuanning Guo, Smruti Mahapatra, Haguy Wolfenson, Serge Ankri

## Abstract

*Entamoeba histolytica* is a pathogenic amoeba that inhabits the human large intestine and causes amoebiasis. *E. histolytica* interacts with both the intestinal microbiota and the metabolites they produce. These bacterial metabolites play a crucial role in shaping the virulence and stress resistance of *E. histolytica*. One such metabolite, indole, is synthesized by bacteria from tryptophan and functions as a key signaling molecule. In this study, we investigated the impact of indole on *E. histolytica* by incubating trophozoites with the metabolite and performing proteomic analyses under various conditions, including trophozoites adapted to indole. Our results show that indole is toxic to *E. histolytica*, with an inhibitory concentration (IC₅₀) of 1.2 mM; however, the parasite can adapt to this concentration. Proteomic analyses reveal that indole-adapted trophozoites display enhanced resistance to oxidative stress (OS), upregulation of cytoskeletal proteins, and increased virulence. These trophozoites exhibit increased F-actin formation, smaller cell size, stronger adhesion to HeLa cells, enhanced migratory capacity, and more effective colonization of the mouse cecum compared to non-adapted trophozoites. Thus, indole exerts a dual effect on *E. histolytica* physiology. While indole is initially toxic to the parasite, adaptation to indole confers enhanced resistance to OS and promotes a more virulent phenotype. This duality underscores the complex role of microbiota-derived metabolites in modulating parasite behavior and highlights indole as a key microbial signal.

**Author summary:** *Entamoeba histolytica* is a unicellular parasite living in the human large intestine and can cause a disease called amoebiasis. Inside the intestine, *E. histolytica* interacts with the microbiota and with the metabolites they produce. One such metabolite is indole, which is produced by gut bacteria from tryptophan and used by human cells and trophozoites. In this study, we explored how indole affects *E. histolytica*. We found that indole is toxic to the parasite at first, but over time, the amoeba can adapt and survive in its presence. Using protein analysis techniques, we discovered that indole-adapted amoebas become more resistant to stress and more virulent. They produce more F-actin, leading to improved mobility, degrade more HeLa cells, and colonize the gut of mice more efficiently than wild-type amoebas (WT). These results show that, while indole initially harms the parasite, it also triggers changes that make it more resistant to stress and more virulent. Our work highlights how metabolites produced by bacteria in the large intestine can influence the behavior of *E. histolytica*.

## Introduction

Amoebiasis, a human disease caused by the protozoan parasite *Entamoeba histolytica*, remains a significant global health challenge, affecting millions of individuals worldwide, particularly in underdeveloped regions with limited access to sanitation and healthcare facilities [1]. This parasite is present in contaminated feces as cysts, a resistant form of the parasite, and once in the host colon, it becomes trophozoites, the infectious form of the parasite. In 10% of the cases, trophozoites become virulent and reach the liver or the lungs, which can lead to the death of the host. This devastating disease is responsible for substantial morbidity and mortality, making it a priority for the development of innovative and effective therapeutic approaches. *E. histolytica* trophozoites are parasitized in the colon, where they are exposed to the gut microbiota, composed of a diverse consortium of microorganisms, that plays a fundamental role in maintaining gut homeostasis and influencing host immune responses [2].

Emerging research has highlighted the intricate interplay between *E. histolytica* and the gut microbiota, underscoring the importance of the gut microenvironment in shaping the pathogenesis of this parasitic infection (for a recent review see [3]). For instance, cultivating *E. histolytica* with *E. coli* O55 can increase *E. histolytica*’s virulence, which is dependent on the contact between the amoeba and bacteria [4]. Furthermore, *E. histolytica* trophozoites showed greater resistance to oxidative stress (OS) after incubation with *E. coli* O55 [5]. Infection with *E. histolytica* can cause dysbiosis marked by a reduction of *Lactobacillus* and *Bacteroides* and an increase in *Bifidobacterium* [6]. Interactions between *E. histolytica* and the gut microbiota are mediated through a dynamic crosstalk, involving the exchange of metabolites, signaling molecules, and host-derived factors. The gut microbiota can influence the expression of specific virulence-related genes in *E. histolytica*, potentially enhancing its pathogenic potential [7]. Conversely, presence of *E. histolytica* modulates the gut microbiota composition, leading to alterations in the microbial community structure and function [6]. Recently, we have discovered that the interaction between *E. histolytica* and gut microbiota is even more complex than previously thought. *E. histolytica* can distinguish between planktonic and biofilm forms of bacteria and degrade these bacterial biofilms through the action of its cysteine proteases (EhCPs) [8].

Although the gut microbiota is increasingly recognized as a modulator of *E. histolytica* behavior, the specific effects of individual microbial metabolites on the parasite’s growth, virulence, and survival remains largely uncharacterized. Among these metabolites, oxaloacetate has emerged as a key player with its unique antioxidant properties, contributing to the parasite’s redox homeostasis [9]. Additionally, our research has shed light on the regulatory role of queuine, a microbial metabolite, in modulating translation processes within the parasite through tRNA modification [10, 11]. Among various bacterial metabolites, indole plays an essential role as a key signaling molecule produced by the gut microbiota through the breakdown of dietary tryptophan that escapes absorption in the small intestine and reaches the colon, where it is converted into indole by the enzyme tryptophanase [12]. Mammalian cells lack the ability to produce indole, making bacterial sources the sole contributors of indole in the human body [13]. While the fecal indole concentration ranges from 0.30 to 6.64 mM, the exact indole concentration in the colon remains unknown but is likely higher [14].

In humans, indole and its derivatives play key roles in maintaining intestinal immune homeostasis [15]. Indole reduces inflammation by suppressing IL-8 secretion, inhibiting NF-κB activation, and promoting IL-10 production [16]. The indole derivative indoleacrylic acid (IA), synthesized by *Bacteroides*, also exerts anti-inflammatory effects via AhR activation [17]. IA further activates the NRF2–ARE pathway, leading to decreased IL-6 and IL-1β secretion, modulation of immune cell function, and regulation of inflammation- and OS-related genes [18]. Thus, indole compounds contribute to anti-inflammatory and antioxidant defenses in the gut.

*E. histolytica* is exposed to indole produced by bacteria from the microbiota. Interestingly, *E. histolytica* also possesses the enzyme tryptophanase, which catalyzes the conversion of tryptophan into indole and pyruvate, an effective process for ATP production [19]. In the presence of 1 mM tryptophan, *E. histolytica* is able to produce 6.8 ± 0.8 µM of indole after 3 days, which is very low compared to the amount of indole that bacteria such as *E. coli* can produce, up to 600 µM in suspension culture [20, 21]. Indole is a well-studied metabolite due to its role in inter-and intra-cellular signaling in bacterial ecosystems. It influences biofilm formation and virulence factors in bacteria [13]. These influences vary between indole-producing and non-producing bacteria. For example, indole enhances biofilm formation in non-indole-producing bacteria such as *Pseudomonas aeruginosa*, whereas it suppresses biofilm formation in indole-producing bacteria like *E. coli* [22].

Our understanding of the effects of indole on parasitic organisms remains limited. In the case of *Cryptosporidium*, higher levels of indole in fecal material have been correlated with reduced severity of diarrhea in patients, suggesting that indole-producing bacteria may impair the parasite’s ability to establish infection in the human intestine, possibly through disruption of mitochondrial functions [23, 24]. Similarly, the indole derivative methyl 6-chloro-1H-indole-3-carboxylate induces mitochondrial dysfunction in the free-living amoebae *Acanthamoeba castellanii* and *Acanthamoeba griffini*, triggering reactive oxygen species (ROS) production and programmed cell death [25]. Beyond protozoa, indole also influences bacterial pathogens: for instance, *Porphyromonas gingivalis*, a major contributor to periodontitis, exhibits decreased virulence when its tryptophanase gene is deleted. This loss of indole production alters its proteome and metabolome, leading to diminished expression of virulence factors, reduced biofilm formation, and impaired host interactions [26]. Given the abundance of indole in the colon and its known effects on other microbes and protozoa, we hypothesize that it may modulate *E. histolytica* physiology and virulence without harming host cells. This study explores the impact of indole on *E. histolytica* trophozoites and the underlying molecular mechanisms.

## Methods

### E. histolytica culture

*E. histolytica* trophozoites of strain HM-1:IMSS (from Prof. Samudrala Gourinath, Jawaharlal Nehru University, New Delhi, India), were grown at 37 °C in 13 × 100 mm in screw-capped Pyrex glass tubes Diamond’s TYI S-33 medium (Johnson and Johnson, Hyclone, USA), supplemented with bovine serum (Biowest, France), to the exponential phase. Trophozoites were harvested from their growth support by incubating the tubes by tapping the glass tubes, followed by centrifugation (Eppendorf centrifuge 5810R, rotor A-4-62) according to a previously reported protocol [23]. Hamster liver-passaged virulent HM-1:IMSS (v-HM1) was cultured in YIMDHA-S medium supplemented with live *Crithidia fasciculata* at 35°C [24] [25].

### HeLa cell cultures

HeLa cells (a kind gift from Dr. Kleinberger, Faculty of Medicine, Technion) were maintained in Dulbecco’s modified Eagle’s medium (DMEM) (Gibco, ThermoFisher Scientific, Inchinnan, Scotland, UK) supplemented with 4 mM l-glutamine, penicillin (100 units ml^−1^), streptomycin (100 μg ml^−1^), and 10% fetal calf serum (Gibco, Thermo Fisher Scientific, Brazil). The media was changed every 2 days. The cultures were grown in 15 by 10 cm plastic tissue culture flasks and maintained in a humidified atmosphere of 5% CO_2_ at 37 °C. The media was changed every 2 days.

### Bacteria strains

All *B. subtilis* strains utilized in this study were derived from the proficient biofilm-forming strain NCIB3610 [26].

### *B. subtilis* Biofilm formation

The biofilm formation was done according to [8]. Briefly, a single colony was isolated from lysogeny broth (LB) plates and grown to mid-logarithmic phase in a 3-ml LB culture, with shaking at 37 °C for 4 h at 200 rpm. For biofilm preparation, cells from the mid-logarithmic phase were diluted 1:10 into a serum-free Diamond’s TYI S-33 medium and grown overnight at 30 °C without shaking. The growth was carried out in 24-well plates, with each well covered by 1 ml of serum-free Diamond’s TYI S-33 agar.

### Biofilm degradation assay

Biofilm degradation essays were performed according to [8]. Briefly, Trophozoites (1 × 10^6^) were incubated on *B. subtilis* biofilm at 37 °C for 3 h for *B. subtilis* and 1 h without shaking. To quantify the extent of biofilm degradation, the GFP signal intensity of each well was compared to the control (biofilm incubated without trophozoites) using ImageJ software.

### IC50

*E. histolytica* trophozoites (1 × 10^6^ cells) in TYI S-33 medium (ThermoFisher, Scientific, Israel) were treated with indole (Sigma-Aldrich, China) at concentrations ranging from 0, 0.5, 1, 2, 3, and 4 mM for 12 hours at 37°C. The viability of the trophozoites was determined by the eosin dye exclusion method [27].

### Growth Rate of *E. histolytica* Trophozoites

A total of 4 × 10^4^ *E. histolytica* trophozoites were grown in a 15 mL tube in TYI-S-33 medium at 37°C. Viable trophozoites were counted after 24 and 48 hours using a previously described protocol [27].

### Adaptation to indole

The concentration of indole in *E. histolytica* trophozoite culture was progressively increased from 0 to 1.2 mM over a period of two months.

### Measurement of cytopathic activity

The rate of destruction of cultured HeLa cell monolayers by *E. histolytica* trophozoites was measured as described in [4]. Briefly*, E. histolytica* trophozoites (3 × 10^5^ per well) were incubated at 37°C with HeLa cell monolayers in 24-well plates for 30 min. The incubation was stopped by placing the plates on ice and washing off the trophozoites with cold PBS containing 1% galactose. The HeLa cells remaining on the tissue culture plate were stained with methylene blue (0.1% in 0.1 M Borate buffer pH 8.7). The dye was extracted from the stained cells using 0.1 M HCl, and its absorbance was measured at 660 nm using a spectrophotometer.

### Detection of intracellular indole by metabolomics

All experiments were performed in quintuplicate: wild type (WT), trophozoites adapted to 1.2 mM indole (ADI), and trophozoites exposed to acute indole (TI, incubated with 1.2 mM indole for 24 hours at 37°C). Trophozoites were harvested by gently tapping the glass tube, followed by centrifugation at 1900 rpm for 3 minutes at room temperature (RT). The cell pellets were dissolved in phosphate-buffered saline (PBS) and transferred to 2 ml homogenizing tubes and centrifuged (1900 rpm for 3 minutes at RT). After removing the PBS, the cells were dissolved in 300 µl of ice-cold metabolite extraction solution (methanol (Merck, 106035), acetonitrile (Merck, 100029) and water at a ratio of 5:3:2). For metabolites extraction, the cell pellet dissolved in extraction buffer was added to CK14 homogenizing tubes containing 1.4 mm ceramic beads (Bertin Corp, P000926-LYSK0-A). Samples were homogenized at 4°C in a Precellys 24 tissue homogenizer. Homogenization conditions were set to three cycles, 30 seconds each, at 6500 rpm with a 30 second gap between each of the cycles to preserve low temperature. Homogenates were centrifuged at 18,000g for 15 minutes at 4°C. The clear supernatants were transferred to glass high-performance liquid chromatography (HPLC) vials (Agilent, 8010-0542) and kept at −80°C until Liquid chromatography-mass–mass spectrometry (LC-MS) analysis.

### Metabolomics Profiling LC-MS analysis

For polar metabolites detection, LC-MS metabolomics analysis was performed as described previously (PMID: 26358905) with some slight changes. ThermoFisher Scientific Vanquish ultra-high-performance liquid chromatography (UHPLC) system coupled to Exploris 240 Orbitrap Mass Spectrometer (ThermoFisher Scientific) was used with a resolution of 60,000 at 200 mass/charge ratio (*m/z*), electrospray ionization, and polarity switching mode to enable both positive and negative ions across a mass range of 67 to 1000 m/z. HPLC setup consisted of ZIC pHILIC column (SeQuant; 150 mm x 2.1 mm, 5µm). 5 µL of biological extracts were injected and the compounds were separated using mobile phase gradient of 15 min, starting at 20% aqueous (20 mM ammonium carbonate (ThermoFisher Scientific, 10785511) adjusted to pH = 9.2, with 0.1% of 25% ammonium hydroxide (ThermoFisher Scientific, 15547049) and 80% acetonitrile. The gradient was terminated with 20% acetonitrile. Flow rate and column temperature were maintained at 0.2 mL/min and 45°C, respectively, for a total run time of 27 minutes. All metabolites were detected using a mass accuracy below 5 ppm. The software Xcalibur (ThermoFisher Scientific) was used for data acquisition.

### Metabolomics data analysis

Skyline V (version 25.1) was used for analysis. Peak areas of metabolites were determined using the exact mass of singly charged ions. The retention time of metabolites was predetermined on the pHILIC column by analyzing an in-house mass spectrometry metabolite library consisting of commercially available standards. Each metabolite’s peak area value was normalized to the value of µg of protein of cells lysates. Metabolite AutoPlotter 2.6 (PMID: 32670572) and Metaboanalyst [28] were used for data visualization.

### Preparation of lysate for proteomics

The samples were precipitated in 80% final concentration of ice-cold acetone overnight at −20°C and washed 3 times with cold 80% acetone. The protein pellets were dissolved in 8.5M Urea, 400 mM ammonium bicarbonate and 10 mM DTT. Protein amount was estimated using Bradford readings. The samples were reduced (60°C for 30 min), modified with 35.2mM iodoacetamide in 100 mM ammonium bicarbonate (room temperature for 30 min in the dark) and digested in 1.5 M Urea, 66 mM ammonium bicarbonate with modified trypsin (Promega), overnight at 37°C in a 1:50 (M/M) enzyme-to-substrate ratio. An additional second trypsinization was done for 4 hours in a 1:100 (M/M) enzyme-to-substrate ratio. The tryptic peptides were desalted using Oasis HLB 96-well µElution Plate (Waters)/homemade C18 stage tip, dried and re-suspended in 0.1% Formic acid in 2% acetonitrile.

### Mass spectrometry analysis for proteomics

The resulting peptides were analyzed by LC-MS/MS using an Exploris 480 mass spectrometer (Thermo) fitted with a capillary HPLC (EV-1000, Evosep One, Denmark). The peptides were loaded onto a 15 cm, ID 150 µm, 1.9-micron Performance column EV1137 (Evosep, Denmark). The peptides were eluted with the built-in Xcalibur 15 SPD (88 min) method. Mass spectrometry was performed in a positive mode using repetitively full MS scan (m/z 380–985, resolution 120,000) followed by DIA scans (10Da isolation windows with 1 m/z overlap, and resolution 30,000).

### Data analysis

The mass spectrometry data was analyzed using the DIA-NN software version 1.9.2 [29] against *Klebsiella pneumoniae* ATCC 700721 MGH, with Min peptide length set to 7, Maximum number of missed cleavages set to 1, Cysteine carbamidomethylation enabled as a fixed modification, and protein N-term acetylation enabled as a variable modification. Peptide- and protein-level false discovery rates (FDRs) were filtered to 1%. Statistical analysis of the identification and quantization results was done using Perseus 1.6.7.0 software [30].

### Resistance to H_2_O_2_

In order to quantify the oxidative stress resistance of trophozoites against WT, WT + I, and ADI strains, trophozoites were exposed to H_2_O_2_ (Supelco, Merck, Darmstadt, Germany). Trophozoites (1 × 10^6^ cells) were incubated with 2.5 mM of H_2_O_2_ for 30 minutes at 37°C. The H_2_O_2_ was then washed away by centrifugation, and the viability of the trophozoites was assessed using the eosin exclusion assay [27].

### Detection of oxidized proteins in trophozoites

The level of oxidized protein was measured with the Oxyblot kit (Protein Carbonyl Assay Kit, Abcam, Cambridge, UK) according to the manufacturer’s protocol. Trophozoites of WT, WT + I, and ADI were exposed to H_2_O_2_ (2.5 mM, for 30 min at 37 °C) and then lysed with 1% Nonidet P-40 (NP-40 in PBS) for 15 min on ice. Equal protein concentrations (20 μg) were processed with the OxyBlot, protein oxidation detection kit. The reagent of the kit, 2,4-dinitrophenylhydrazine (DNPH) is binding to carbonyl group acquired by protein during oxidation. The DNPH is then immunodetected and provides quantitative results on the amount of oxidized protein.

### Assessment of protein synthesis through surface sensing of translation (SUnSET)

The SUnSet was performed according to [31]. Briefly, treated trophozoites (2 × 10^6^) were incubated with 10 μg/mL puromycin (Merck, Rehovot, Israel), a structural analog of tyrosyltRNA, for 20 min at 37°C. Following this, the trophozoites were lysed using a lysis buffer (10% NP-40, 0.5M E64 (ChemScene, New Jersey, USA), 100 mM PMSF in PBS). The whole proteins were separated on a 10% SDS-PAGE in SDS-PAGE running buffer and subsequently electrotransferred to a nitrocellulose membrane in protein transfer buffer. Loading equivalency was determined by Ponceau-S (Merck, Rehovot, Israel) staining of the membrane before immunostaining. Puromycin incorporation was detected by immunoblotting using a 1:1000 dilution of monoclonal puromycin antibody (12D10 clone, Merck Millipore, Rosh-Ha’ayin, Israel). Following incubation with the primary antibody, the blots were treated with a 1:5000 dilution of secondary antibody (Jackson ImmunoResearch, West Grove, PA, USA) for 1 h at room temperature, then developed using enhanced chemiluminescence (WesternBright^TM^ ECL, Advansta, CA, USA) and photographed with Fusion FX7 Edge Spectra. Protein synthesis quantification was determined from the intensity of the immunoreactive blots (densitometry) using Fiji software [32].

### Actin staining

A total of 1 × 10^6^ *E. histolytica* trophozoites (WT, WT + I, and ADI) were harvested. Trophozoites (1.5 × 10^5^ trophozoites/mL) were then suspended in TYI medium without serum at 37 °C with and without indole (2.5 mM in DMSO) and transferred onto Ibidi µ-Slide 8 Well plate (80826, ibidi GmbH, Gräfelfing, Germany) and allowed to adhere to the surface during a 1 h incubation at 37 °C. Subsequently, the attached trophozoites were fixed with prewarmed 37 °C 4% paraformaldehyde (PFA, Electron Microscopy Sciences, Hatfield, PA, USA) for 30 min at room temperature. Following fixation, trophozoites underwent three five-minute washes with mild agitation in 0.1% Triton X-100/0.1% Tween 20 in PBS (PBSTT). To ensure complete permeabilization, they were subsequently incubated in PBSTT for an extra five minutes at room temperature. The cells were then quenched with 50 mM NH_4_Cl in PBS for 30 min at room temperature, followed by three washes and blocking with 5% bovine serum albumin (BSA, MP Biomedicals, Solon, OH, USA) in PBSTT for 1 h at room temperature. The next day, the samples underwent three washes PBSTT, followed by a 4 h incubation at 4 °C with a 1:250 dilution of

Alexa Fluor 488 (Jackson ImmunoResearch, West Grove, PA, USA) and 1:500 dilution of Phalloidin-iFluor 594 Reagent (#ab176757, Abcam, Cambridge, UK) in 1% BSA in PBSTT. Finally, the samples were washed twice with PBSTT and two more times with PBS and mounted with SlowFade Gold Antifade Mountant (#S36937, Invitrogen). Imaging was performed using Zeiss LSM800 confocal microscope with a 20x objective. To ensure consistency in confocal imaging, all images were acquired using uniform staining protocols and identical microscope acquisition settings.

### Quantification of fluorescent intensity

To ensure consistency in planar confocal imaging of fixed samples, all images were acquired using uniform staining protocols and identical microscope acquisition settings. Cell segmentation was performed using the Cellpose-SAM model, which provided precise masks delineating individual cells (https://huggingface.co/spaces/mouseland/cellpose)[33]. These segmentation masks were subsequently imported into Fiji for quantitative analysis of fluorescence intensity. The Imaris software was used to analyze 3D images, which automatically detected and counted the trophozoites stained by Phalloidin. The intensity of Phalloidin fluorescence and the volume of trophozoites were assessed using the Imaris software.

### Transwell migration assays

Transwell migration assays were performed in 5 mm transwell inserts (SPL Insert Hanging, 8 µm pore size, Gyeonggi-do, Korea) suspended by the outer rim within individual wells of 24-well plates using a previously described protocol [34]. Trophozoites WT, WT + I and ADI were washed one time in Diamond’s TYI-S-33 medium without serum, and 500 µl aliquots of cells at a concentration of 3×10^5^ trophozoites/ml in serum-free medium were loaded into the upper chamber of the transwell inserts, which were then placed in anaerobic bags (Mitsubishi Gas Chemical Company, Inc., Tokyo, Japan), and incubated at 37°C for three hours. At the end of the incubation, the inserts and media were removed, and trophozoite migration was assessed by counting the number of trophozoites attached to the bottom of the well. To facilitate detachment, the plate was placed on ice for 10 minutes. The cells were then counted using the eosin assay.

### Intra-cecum injection in mice

The liver-passaged v-HM1 was maintained by serial passage through the mouse cecum. Trophozoites isolated from the mouse cecum were monoxenically cultured in the YIMDHA-S medium supplemented with *C. fasciculata* [35] with a stepwise increase in indole concentration. After 5 weeks, the indole concentration was increased to 1.2 mM. Six-week-old male C57BL/6NCrSlc mice were purchased from Japan SLC (Inc. Shizuoka, Japan). Mice were anesthetized with isoflurane and surgically injected 1 × 10^6^ cecum-passed v-HM1 in 200 µL of culture medium into the proximal and apical site of the cecum [35, 36]. Prior to formal challenge, indole susceptible v-HM1 and 1.2 mM indole-attenuated v-HM1 were passed through the cecum of mice for 24 h to allow recovery and initiation of axenization. Cecum-recovered v-HM1 strains were challenged again to mouse cecum within 2 weeks of axenic culture.

### Detection and quantification of trophozoites in mouse stool

Body weight was monitored daily for 24 hours after the challenge. After seven-days post infection, DNA was extracted from 3 - 5 stool particles using the QIAcube robotic device and the QIAamp Fast DNA stool Mini Kit (Qiagen, Tokyo, Japan). Infection was assessed by PCR targeting the 18S ribosomal RNA gene of *E. histolytica* (GenBank: AB282658). Primer sequences used in this study were as follows; EntaF2 (5’-ATCCATGATCGCTATAAGATGCACGAGAG-3’) and EhR-4 (5’-CCATAAACTCAAGATTTCTCTTTAAGTTCTGAACAA-3’). To quantify the number of trophozoites in stool, real-time quantitative PCR was performed by using Fast SYBR green master mix (Thermo Fisher Scientific, USA) in the StepOne Plus real-time PCR system (Applied Biosystems, USA) [10].

### Ethics statement and animal experiment

The animal experiments were approved by the Institutional Animal Care and Use Committee (approval number: 123041) and were conducted at the AAALAC-accredited National Institute of Infectious Diseases, JIHS, Japan. At the end of the experiments, the animals were euthanized by cervical dislocation under anesthesia with isoflurane.

### Statistical analysis

Statistical analyses were performed with GraphPad Prism 9.0 (GraphPad 234 Software, Inc., San Diego, CA). The test used for pair-wise experiments was an unpaired t-test. The P-value were, *p < 0.05, **p < 0.01, ***p < 0.001, and ****p < 0.0001. Otherwise, the test used was a multiple comparison test (ANOVA) as indicated in the legends of each figure. Statistical tests are mentioned in the indicated legends of the figures.

## Results

### *E. histolytica* viability and adaptation to indole exposure

The impact of indole on *E. histolytica* in the human colon on the parasite’s physiology remains largely unexplored. To address this, we evaluated the cytotoxicity of indole on *E. histolytica* trophozoites. The trophozoites were exposed to varying concentrations of indole for 24 hours at 37°C. Our findings indicate that indole affects the viability of *E. histolytica*, with an observed IC_50_ of 1.2 ± 0.073 mM after 24 hours. Additionally, previous researches show that *E. histolytica* can adapt to various physiological challenges and toxic conditions, including glucose starvation, exposure to auranofin, and metronidazole treatment [37–39]. This adaptability suggests that *E. histolytica* might also develop responses to indole-induced stress. To explore this, we investigated the parasite’s capacity to adapt to indole. Trophozoites were progressively exposed to increasing concentrations of indole until reaching 1.2 mM (trophozoites adapted to 1.2 mM indole: ADI strain). At this concentration of indole, the doubling time of the adapted ADI strain takes 9 hours, compared to 7.5 hours for the control WT strain cultivated without indole. The IC_50_ of the ADI strain increased to 3-fold (3.5 mM ± 0.07 mM) that of the control WT strain.

To understand how indole could cause this toxic effect on trophozoites, the capacity of the indole to penetrate trophozoites was investigated. Indole acts as an intercellular signal in microbial communities [12], as it can cross bacterial lipid membranes, such as those in *E. coli*, without requiring a protein-based transport system, which suggests that it could potentially diffuse through membranes in other biological kingdoms as well [40]. Therefore, we hypothesize that indole may similarly diffuse across the membrane of *E. histolytica* trophozoites and accumulate in their cytoplasm, initiating a process that ultimately leads to their death. To test this hypothesis, we quantified indole levels in *E. histolytica* trophozoites using metabolomics analysis under three different conditions: wild-type trophozoites not exposed to indole (WT), wild-type trophozoites exposed to 1.2 mM indole for 24 h (WT + I), and trophozoites adapted to 1.2 mM indole (ADI). The intracellular indole levels were significantly higher in both WT + I and ADI trophozoites compared to WT, indicating that indole was incorporated into *E. histolytica* cytoplasm (Fig. 1). A low indole level was detected in WT, indicating an endogenous indole production from tryptophan through the enzyme tryptophanase [16].

**Fig 1.**
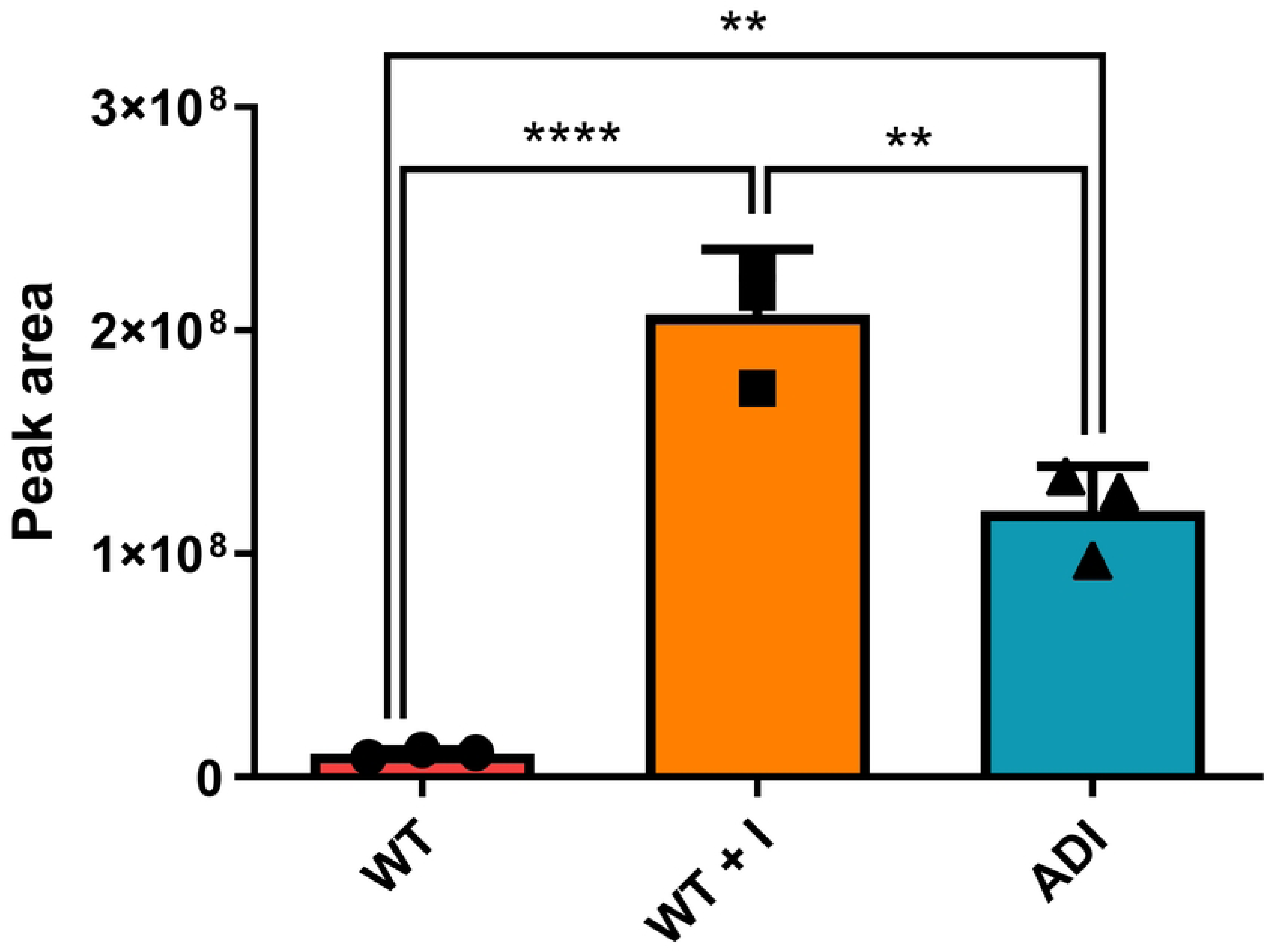
Intracellular level of indole in WT, TI, and ADI. Indole concentration in lysate from WT, TI, and ADI trophozoites was measured by liquid chromatography-mass spectrometry (LC-MS). Statistical analysis was performed using a one-way ANOVA, with significance denoted by * indicating a p-value of <0.05, ** for p-value of <0.01, *** for p-value of <0.001. The data represent the average results obtained from three biological replicates.

### Protein expression changes in *E. histolytica* in response to indole

To investigate the effect of indole on *E. histolytica*, we conducted a proteomic analysis comparing protein abundance in WT + I and ADI, and WT trophozoites (S1 Table). The heatmap clearly shows a distinction in proteomic profiles, highlighting significant differences between, on one side, WT and WT + I, and the other side, ADI trophozoites (Fig. 2A). These findings suggest that trophozoites have a different proteomic response after adaptation to indole. To further explore the molecular basis of this adaptation, we analyzed differential protein expression and categorized the affected proteins using PANTHER, a bioinformatics tool that classifies proteins into functional classes.

**Figure 2:**
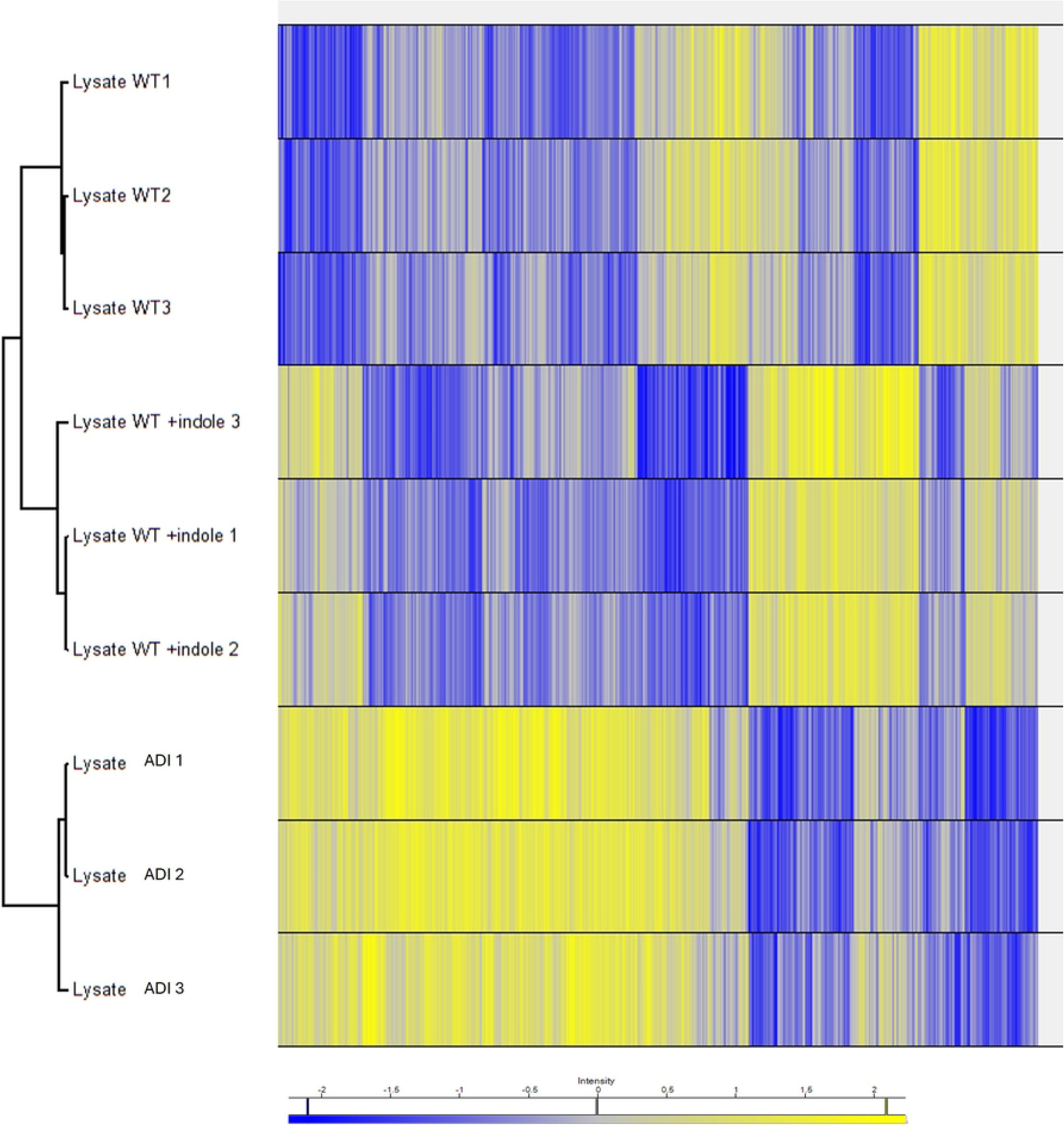
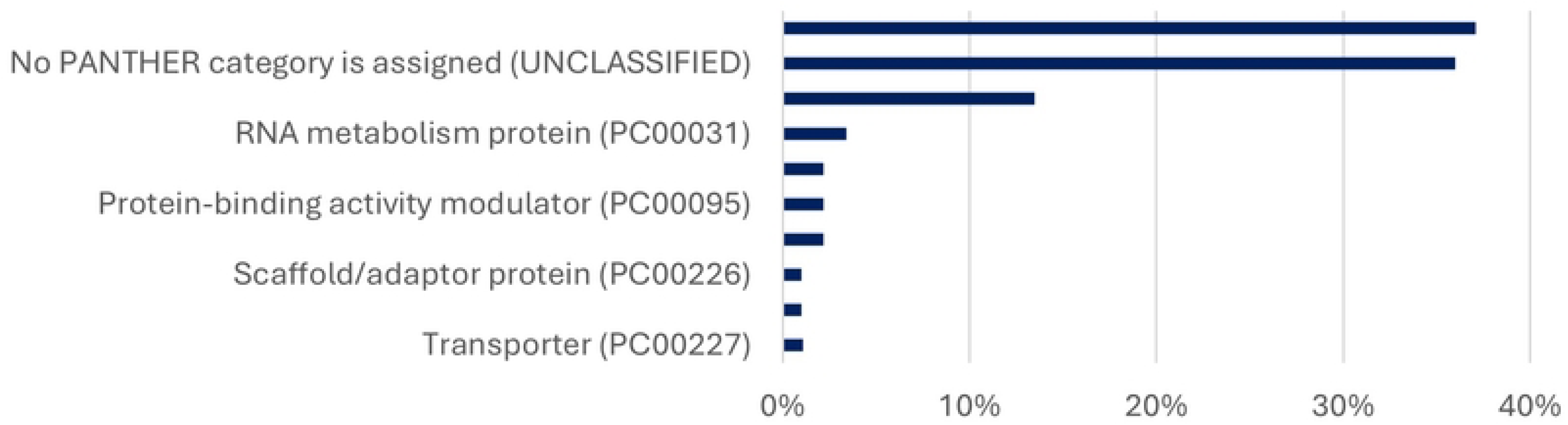
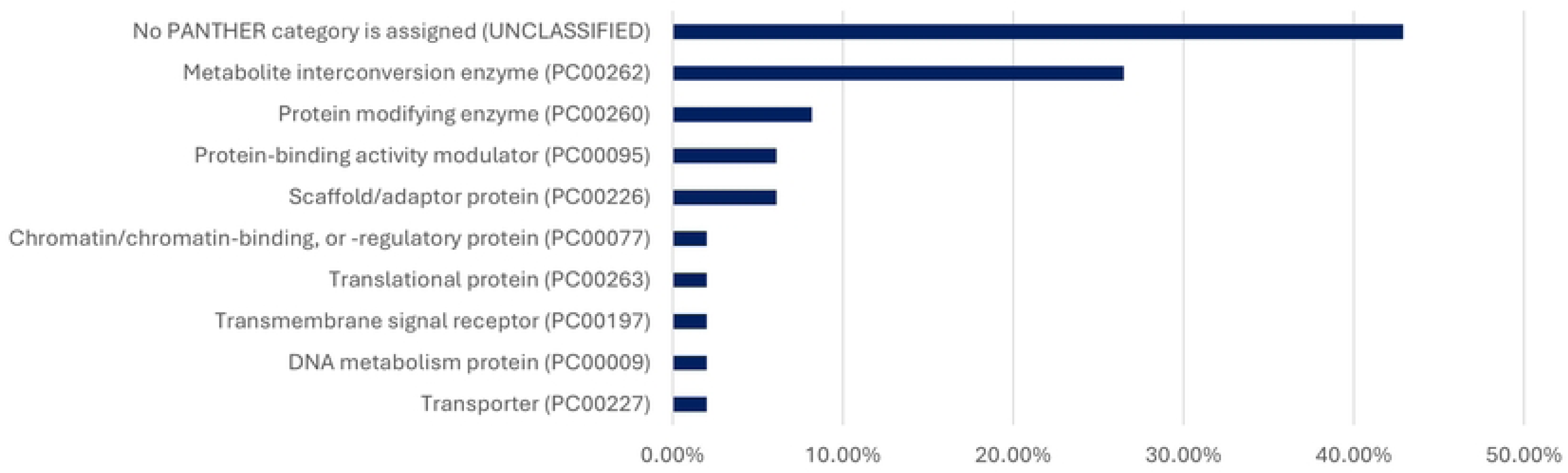
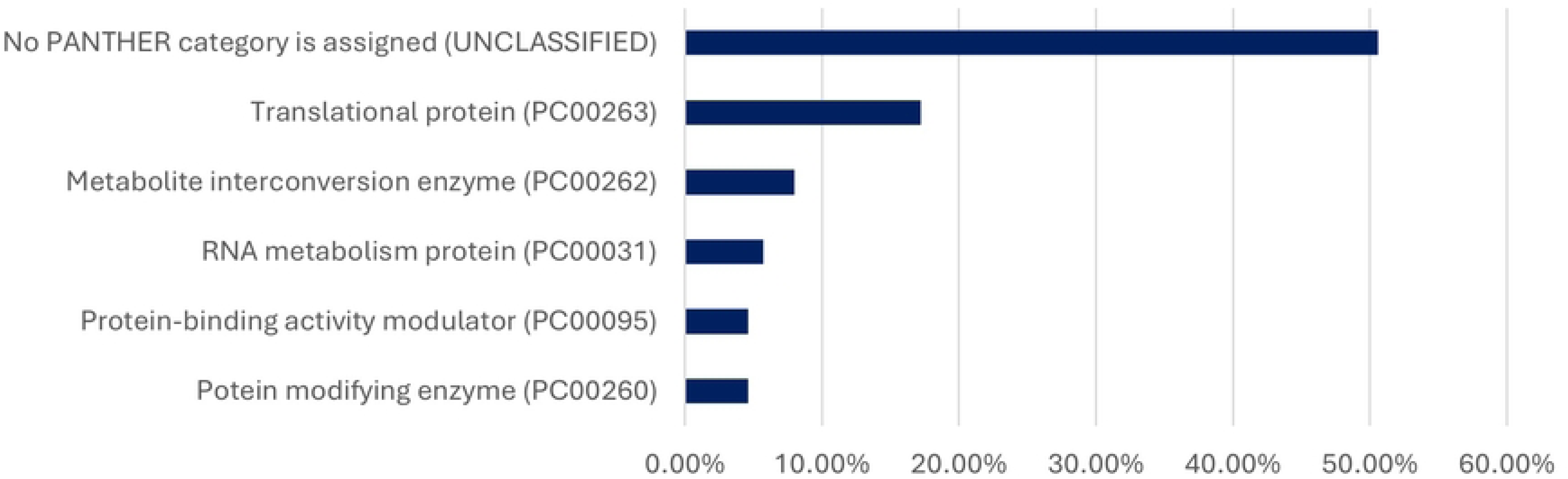
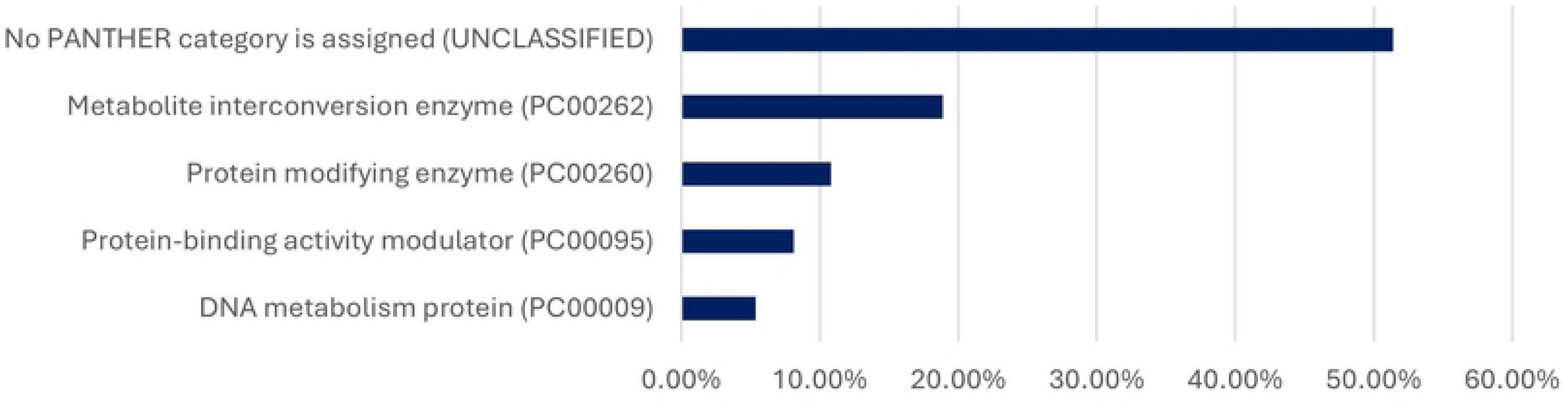
Protein classes enriched in WT, TI, and ADI trophozoites. (A) Heatmap of the proteomics results. (B) Upregulated in ADI compared to WT+I, (C) Downregulated in ADI compared to WT+I, (D) Upregulated in ADI compared to WT, (E) Downregulated in ADI compared to WT.

Upregulated proteins in ADI compared to WT include translational protein (PC00263) and ribosomal protein (PC00202), indicating an upregulation of protein translation. Cytoskeleton-related proteins are also upregulated in ADI compared to WT, suggesting a more dynamic cytoskeleton. For instance, Cytoskeletal proteins such as Actin putative (EHI_043640), protein-binding activity modulator (PC00095), and Rho guanine nucleotide exchange factor (EHI_100140). Virulence factors or virulence-related proteins are found to be both upregulated and downregulated in ADI compared to WT. Protein modifying enzymes (PC00260), where cysteine proteinase is the perfect example; four cysteine proteases (EHI_180170, EHI_010580, and EHI_039310) are upregulated, while CP1 is downregulated in ADI compared to WT. Eukaryotic translation initiation factor 5A (EHI_151540) and Galactose binding lectin 35 kDa subunit (EHI_027800) are upregulated in ADI compared to WT. However, pore-forming peptide amoeba pore B (EHI_194540) is downregulated (Fig. 2D and E, and Fig 3C and D, S1 Table).

**Figure 3:**
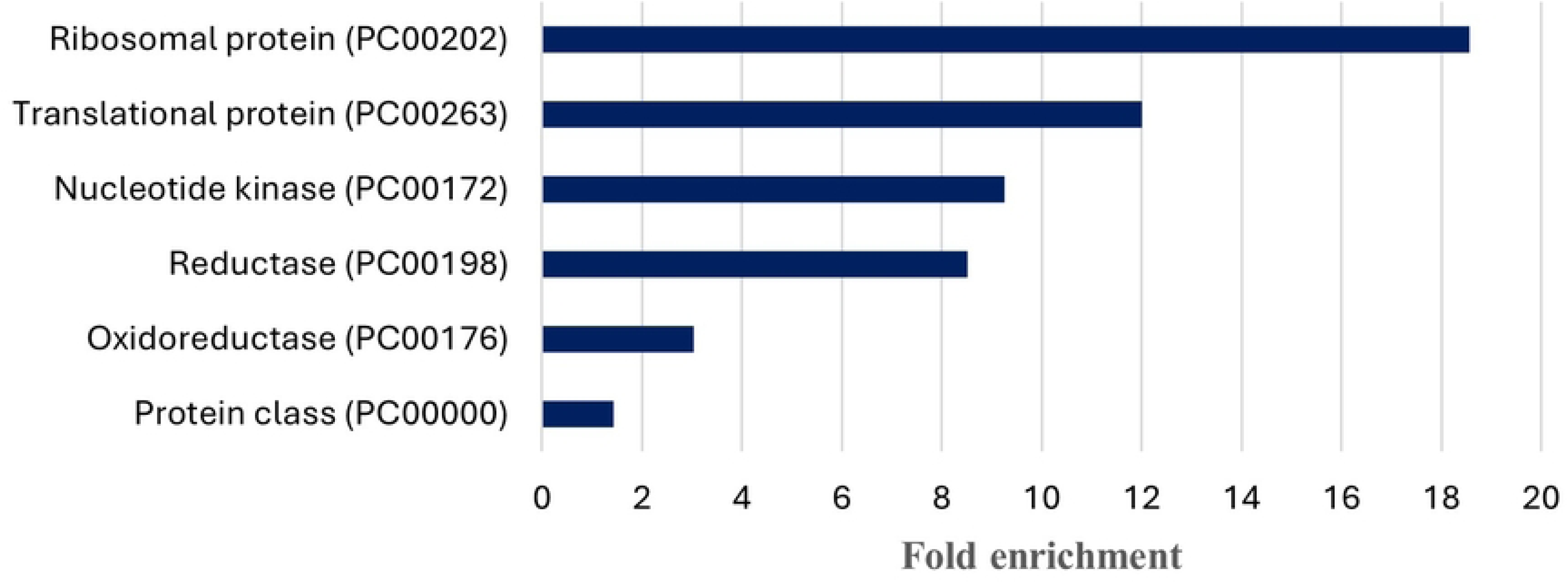
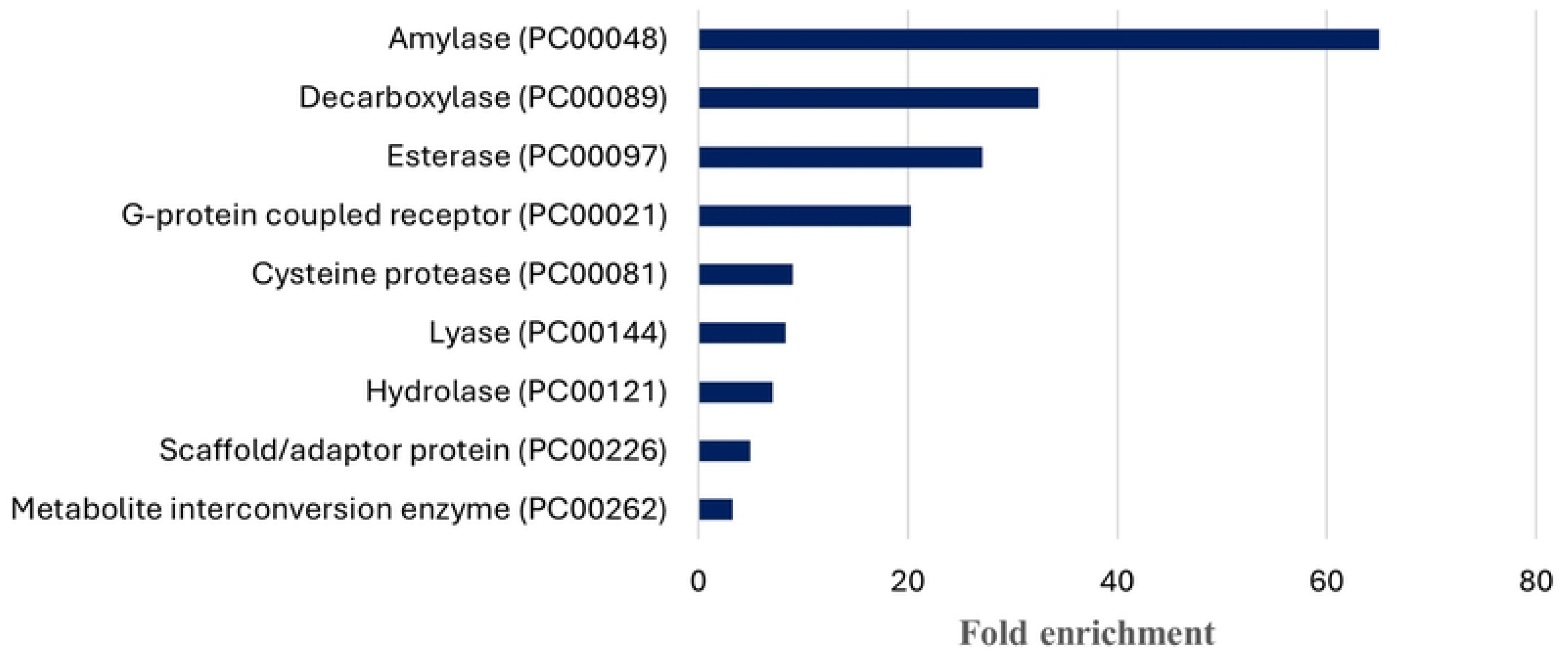
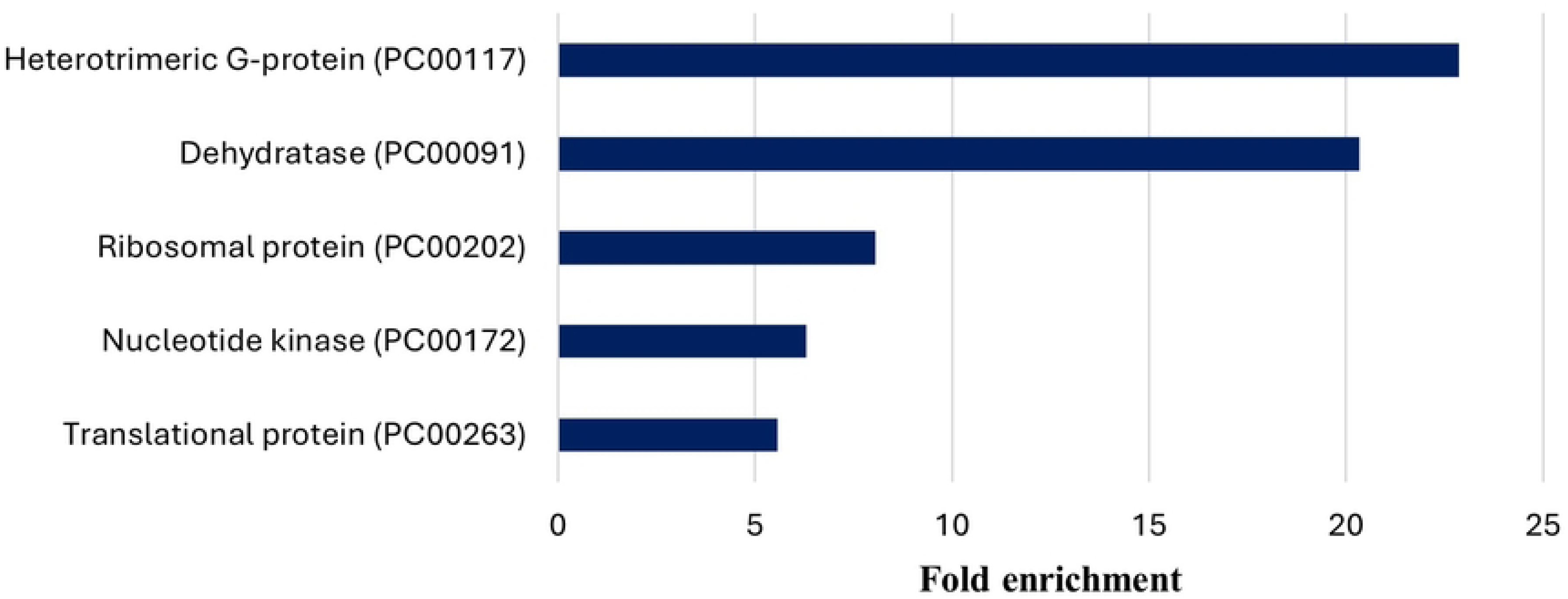
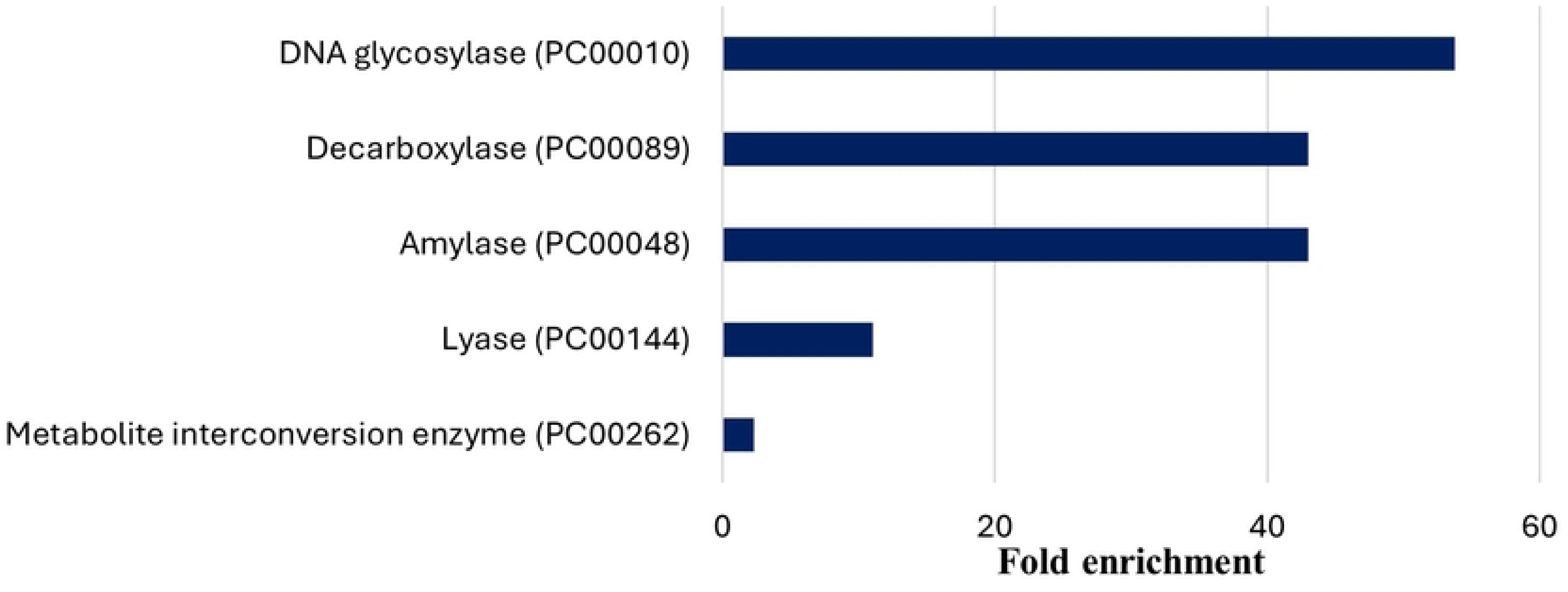
Fold enrichment for protein classes found in WT, TI, and ADI trophozoites. (A) Upregulated in ADI compared to WT+I, (B) Downregulated in ADI compared to WT+I, (C) Upregulated in ADI compared to WT, (D) Downregulated in ADI compared to WT.

ADI compared to WT + I, shows similar upregulated proteins to ADI compared to WT. For instance, the upregulation of translational protein (PC00263) such as 40S ribosomal protein (EHI_105180), or 60S ribosomal protein (EHI_152570), cytoskeletal protein (PC00085) such as Actin (EHI_043640) and protein-binding activity modulator (PC00095) such as Rho guanine nucleotide exchange factor, (EHI_110980) (Fig 2 B and Fig 3A). In addition, ADI, compared to WT + I, presents an upregulation of oxidoreductase (PC00176) such as malate dehydrogenase (EHI_030810) and chaperone (PC00072), like heat shock protein 70, putative (EHI_150770), suggesting an increased resistance of ADI to oxidative stress. The virulence factor Galactose binding lectin 35 kDa subunit (EHI_027800) is also upregulated in ADI compared to WT + I, (Fig. 2B and Fig 3A). However, other virulence factors are downregulated in ADI compared to WT + I, like cysteine proteinase (EHI_151440, EHI_050570, EHI_074180) or pore-forming peptide amoebapore A (EHI_159480), pore-forming peptide amoebapore B (EHI_194540), sending a mixed message about the increase or decrease of virulence of the ADI trophozoites (Fig. 2C and Fig 3B). Even cytoskeleton proteins, heat shock proteins, or virulence-related proteins are all upregulated in AI compared to both WT and WT + I. These proteins’ fold change is more important in ADI compared to WT + I, showing a specific response to indole after adaptation of the trophozoites (Fig. 2A, and S1 Table).

### ADI trophozoites are more resistant to oxidative stress (OS)

Proteomics data indicate an upregulation of oxidoreductase proteins, including peroxiredoxin [41], in ADI compared to WT + I, suggesting that redox regulation plays a role in adaptation to indole. Among the oxidoreductase proteins, Glyceraldehyde-3-phosphate dehydrogenase (GAPDH) enzymes (EHI_060860), or Glutamate synthase beta subunit (EHI_110520), are upregulated in ADI relative to WT + I (Fig. 3C and S1 Table). Based on these findings, we hypothesized that trophozoites adapted to indole exhibit increased resistance to OS. To test this, we induced OS by incubating trophozoites (WT, WI+I, and ADI) with H_2_O_2_ (2.5 mM H₂O₂ for 30 minutes) and quantified the cell survival after exposure to OS. Cell viability was assessed using the eosin assay, revealing that ADI trophozoites displayed significantly higher resistance to H₂O₂ compared to both WT and WT + I (Fig 4A). We evaluated whether the altered viability of ADI trophozoites was associated with their ability to protect proteins from oxidation by quantifying protein carbonylation, a hallmark of oxidative damage, under basal conditions and upon exposure to OS (Fig 4B). The results indicated no significant difference in the level of oxidized proteins between WT, WT + I, or ADI when they are exposed to oxidative stress (Fig 4C).

**Figure 4:**
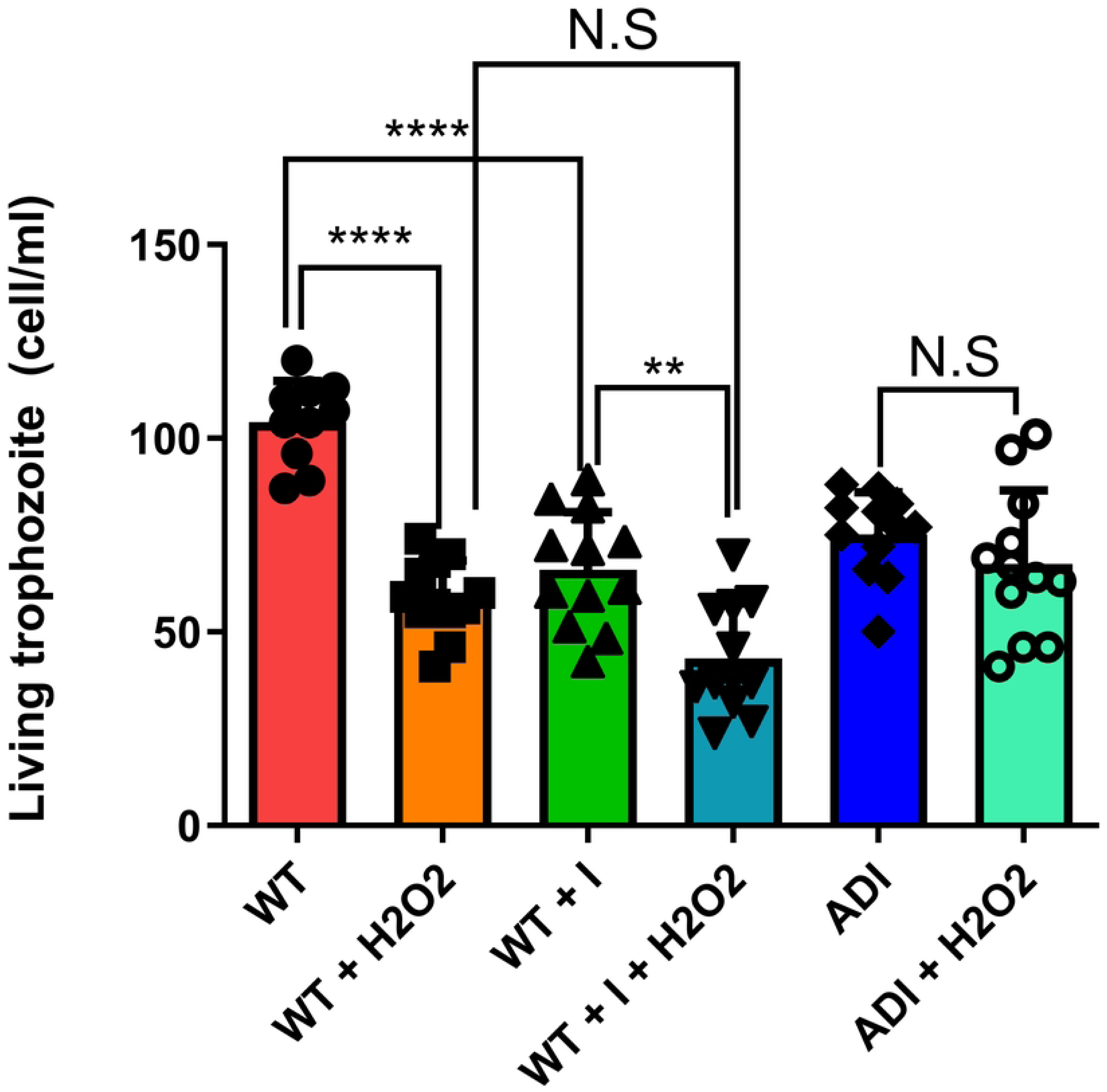
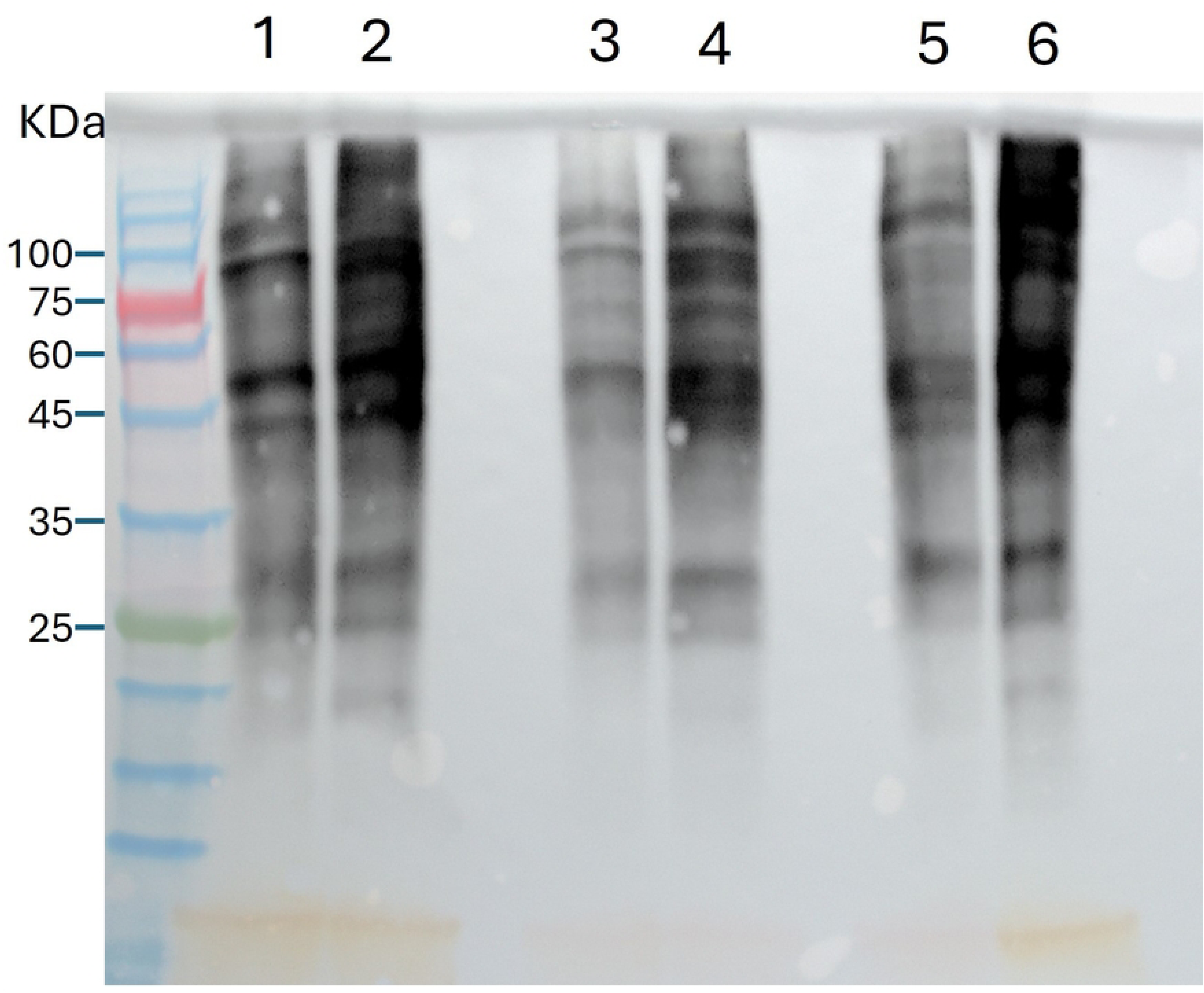
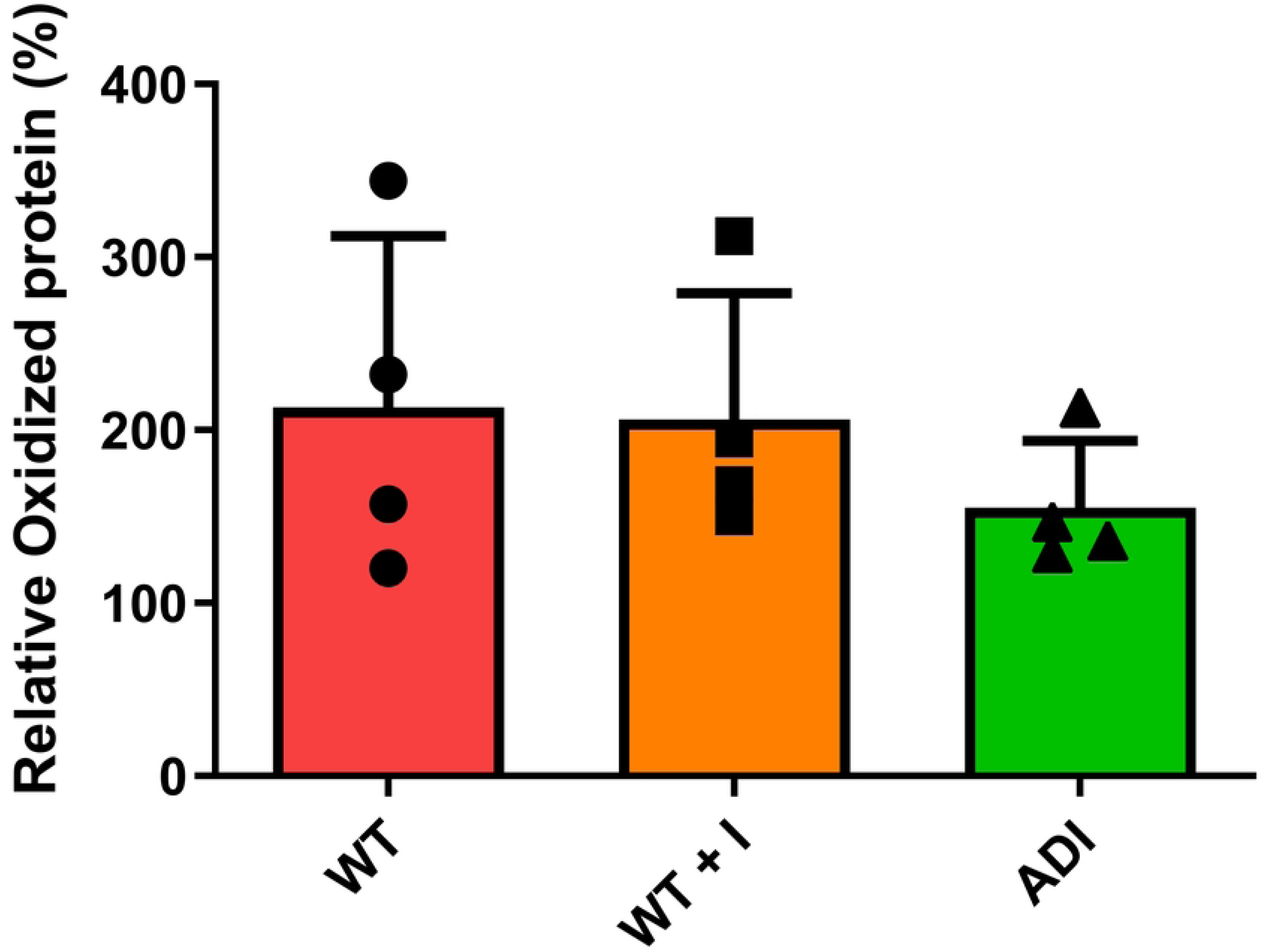
Response of trophozoites to exposure to H_2_O_2_ and quantification of oxidized proteins. (A) Trophozoites (WT, WT+I, and ADI) were incubated with H_2_O_2_ (2.5 mM) for 30 minutes at 37°C. The trophozoite’s survival was quantified with an eosin assay. (B) Gel showing the detection of carbonyl group, indicating oxidized proteins in trophozoites using the OxyBlot method, in the presence or absence of H_2_0_2,_ (1) WT, (2) WT + H_2_O_2_, (3) WT+I, (4) WT+I + H_2_O_2_, (5) ADI and (6) ADI+ H_2_O_2_. (C) Graphic representation of the data from the Oxyblot analysis. The graph shows a ratio between with H2O2/without H_2_O_2_. One-way ANOVA test was performed, ***p < 0.001 and ****p < 0.0001. Data represent the average results from three (A) and four (B) biological replicates.

### Effect of indole on protein translation

Proteins involved in translation are upregulated in ADI compared to WT and WT+I. Previous studies have shown that protein translation levels increase in response to other stressors, such as heat shock [42]. Notably, the translation initiation factor 5A (eIF5A1, EHI_151540) is significantly upregulated in ADI (fold change of 64.3) and is known to play a key role in elongation, termination, and stimulation of peptide bond formation [43]. Additionally, multiple ribosomal subunits show increased expression in ADI compared to WT. These include small (40S) ribosomal proteins (S23, S13, S28, S26, S17, S6, and S18), as well as large (60S) ribosomal proteins (L31, L38, L24, L35a, and L11). Heterotrimeric G-proteins, which are membrane-associated proteins that couple with seven transmembrane receptors to transmit intracellular signals, are also upregulated in ADI. These findings suggest that ADI trophozoites have developed a more efficient translational response as an adaptive mechanism to indole exposure. To test this hypothesis, we have quantified protein synthesis in WT, WT + I, and ADI using the SUrrogate marker for SEquential Translation (SUnSET) technique, which measures protein synthesis through the incorporation of puromycin into nascent polypeptides. The results show that, although there is a trend showing an increase in newly synthesized protein in ADI, the difference is not significant (Fig. 5A&B).

**Figure 5:**
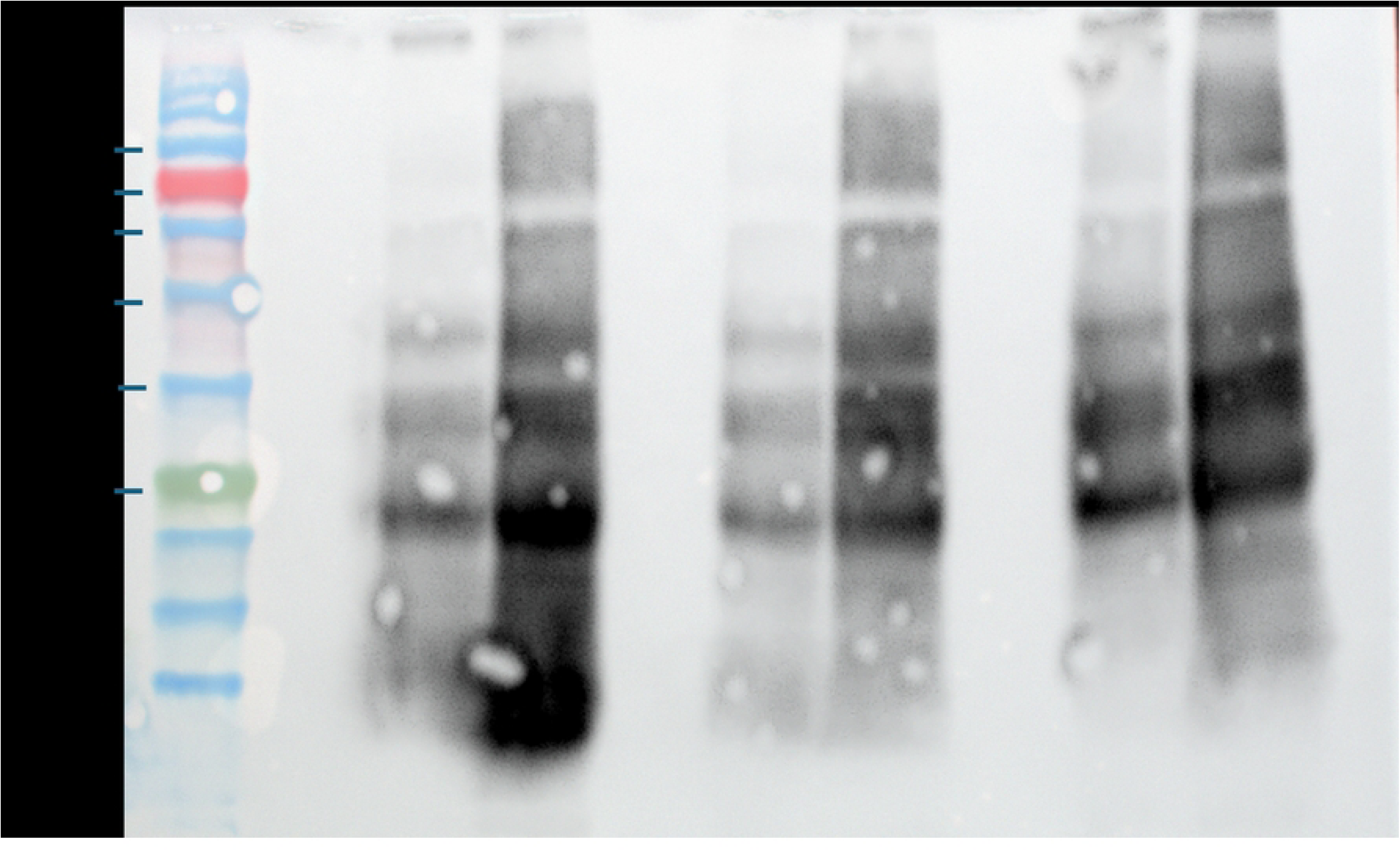
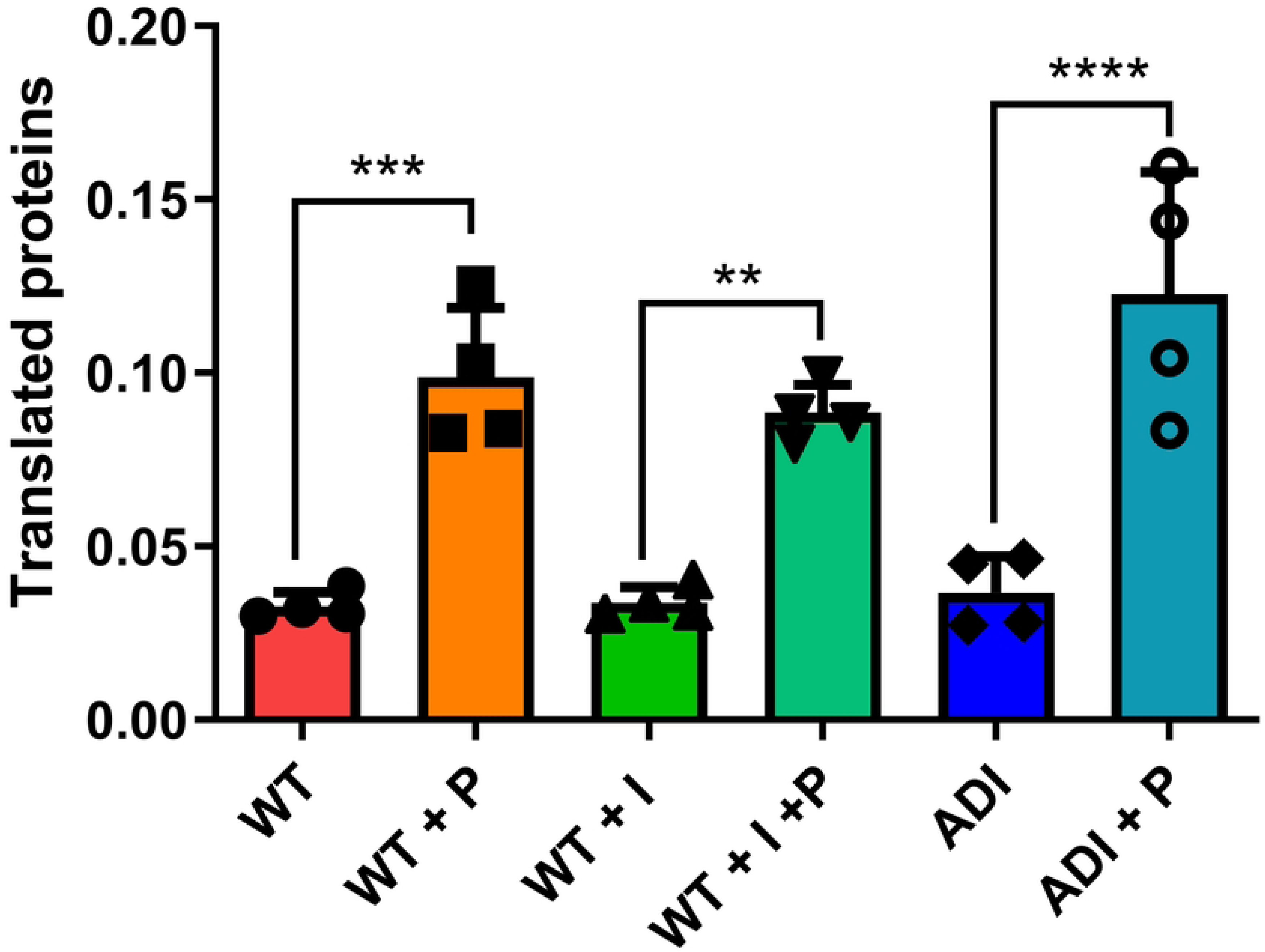
Indole does not affect the synthesis of newly translated proteins in trophozoites (WT, WT+I, and ADI). (A) Sunset Gel showing the detection of newly translated proteins with puromycin incorporation. (1) WT, (2) WT + cycloheximide (100 µg/mL) + puromycin, (3) WT + puromycin, (4) WT+I, (5) WT + I + cycloheximide + puromycin, (6) WT + I + puromycin, (7) ADI, (8)ADI + cycloheximide + puromycin, (9) ADI + cycloheximide + puromycin. (B) Graphic representation of the data from the Sunset analysis. One-way ANOVA test was performed, ***p < 0.001 and ****p < 0.0001. Data represent the average results from four biological replicates.

### Increased level of filamentous actin (F-actin) and trophozoite motility in ADI trophozoites

An actin protein, EHI_043640, is upregulated in ADI compared to both WT and WT + I (S1 Table). Interestingly, this upregulated protein is a truncated version of an actin protein (EHI_182900), and it is not clear if this truncated version is a functional protein [44]. Actin plays a crucial role in *E. histolytica*’s ability to migrate, divide, exert cytotoxicity, and in the phagocytosis of human cells [44], particularly in the form of F-actin, which regulates the assembly and disassembly of microfilaments in amoeba [44]. Other cytoskeleton-related proteins were found upregulated, like α-actinins (EHI_161200), involved in amoeboid motility and phagocytosis, as well as regulators of actin dynamics [45], such as RhoGAPs and RhoGEFs, which collectively support active cytoskeletal remodeling in ADI.

To determine if the increase in cytoskeletal proteins observed in ADI has an effect on F-actin formation, we compared the level of F-actin in WT, WT + I, and ADI trophozoites (Fig 6A and S1A appendix). First, two-dimensional microscopy images and three-dimensional z-stack images were taken with fluorescent confocal microscopy and were analyzed, respectively, with the Cellpose-SAM model coupled with Fiji and Imaris software. Results indicate that F-actin formation measured by fluorescent intensity was increased in ADI compared to WT and WT + I (Fig 6C). The amount of F-actin was also increased in WT + I relative to WT (Fig. 6C). Interestingly, ADI are smaller in size (lower cell area and cell volume) than WT or WT + I (Fig 6C and S1B appendix).

**Figure 6:**
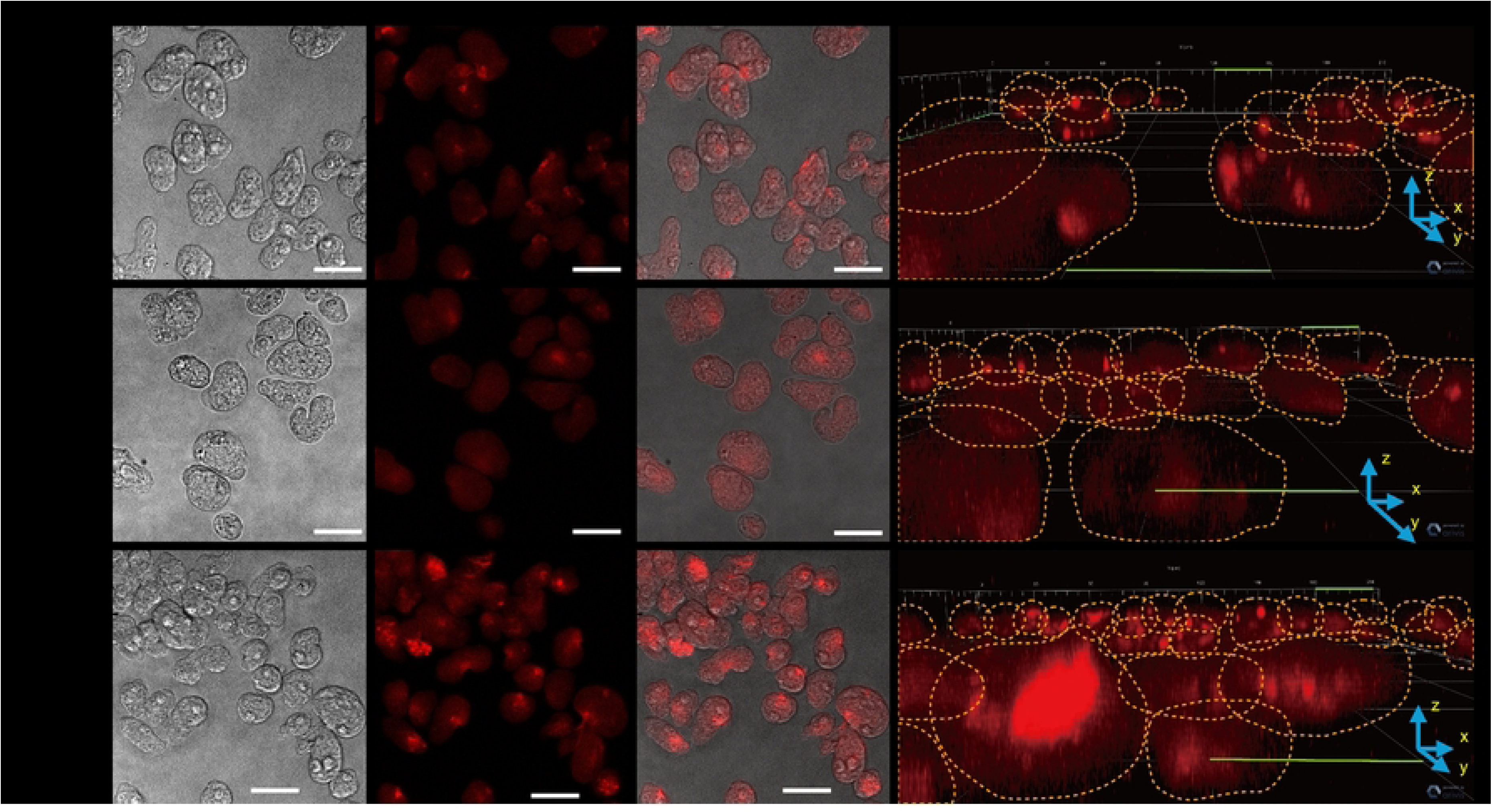
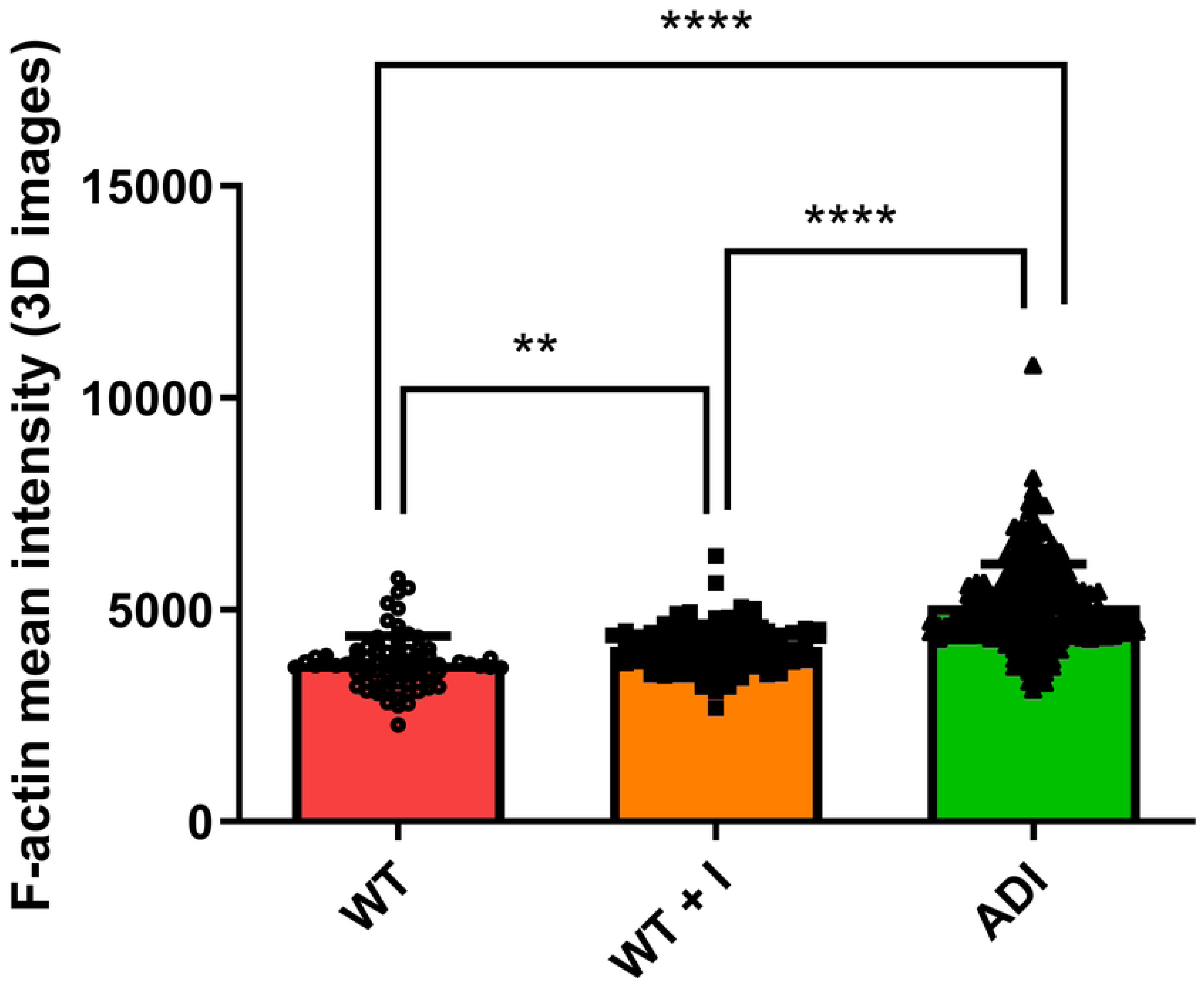
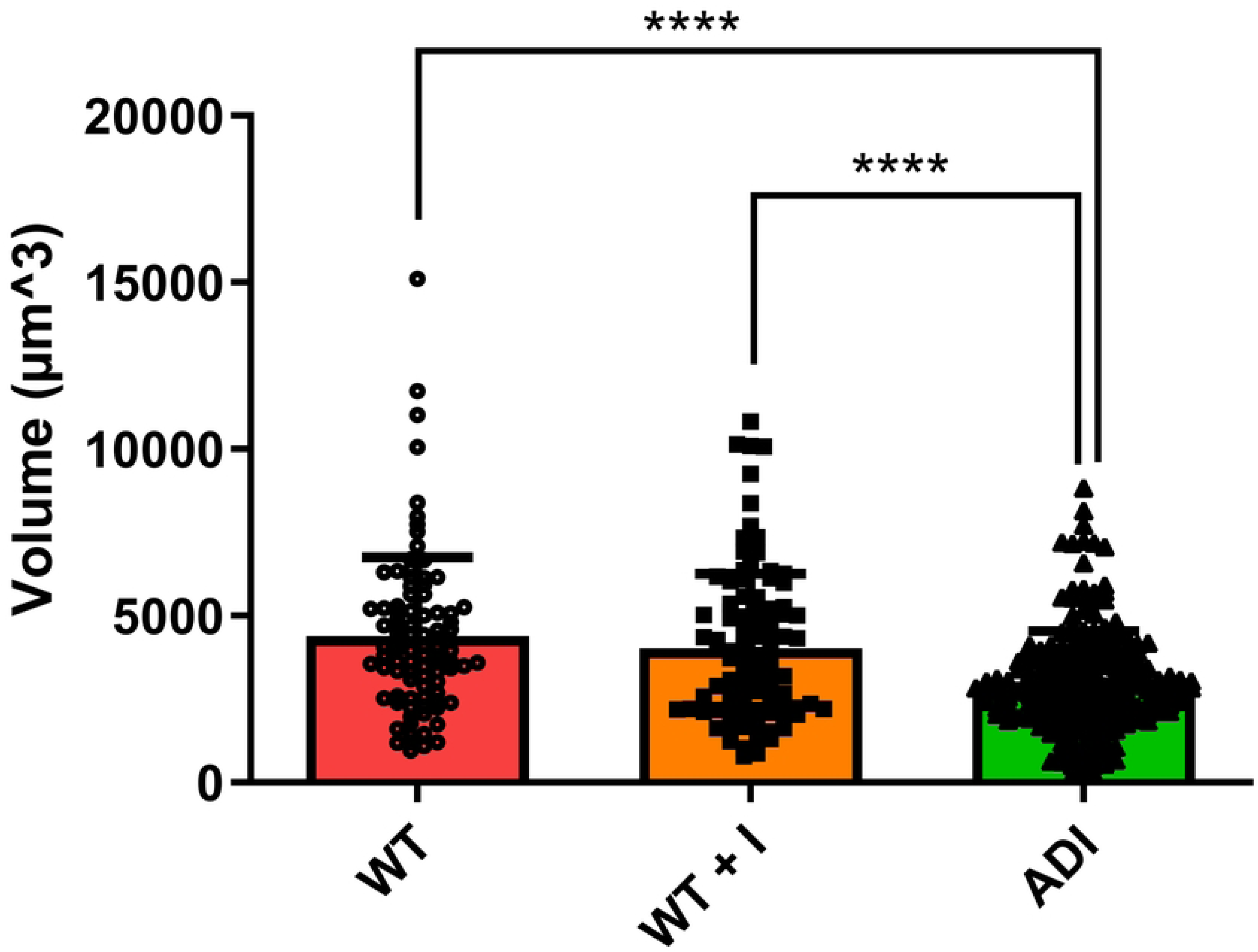
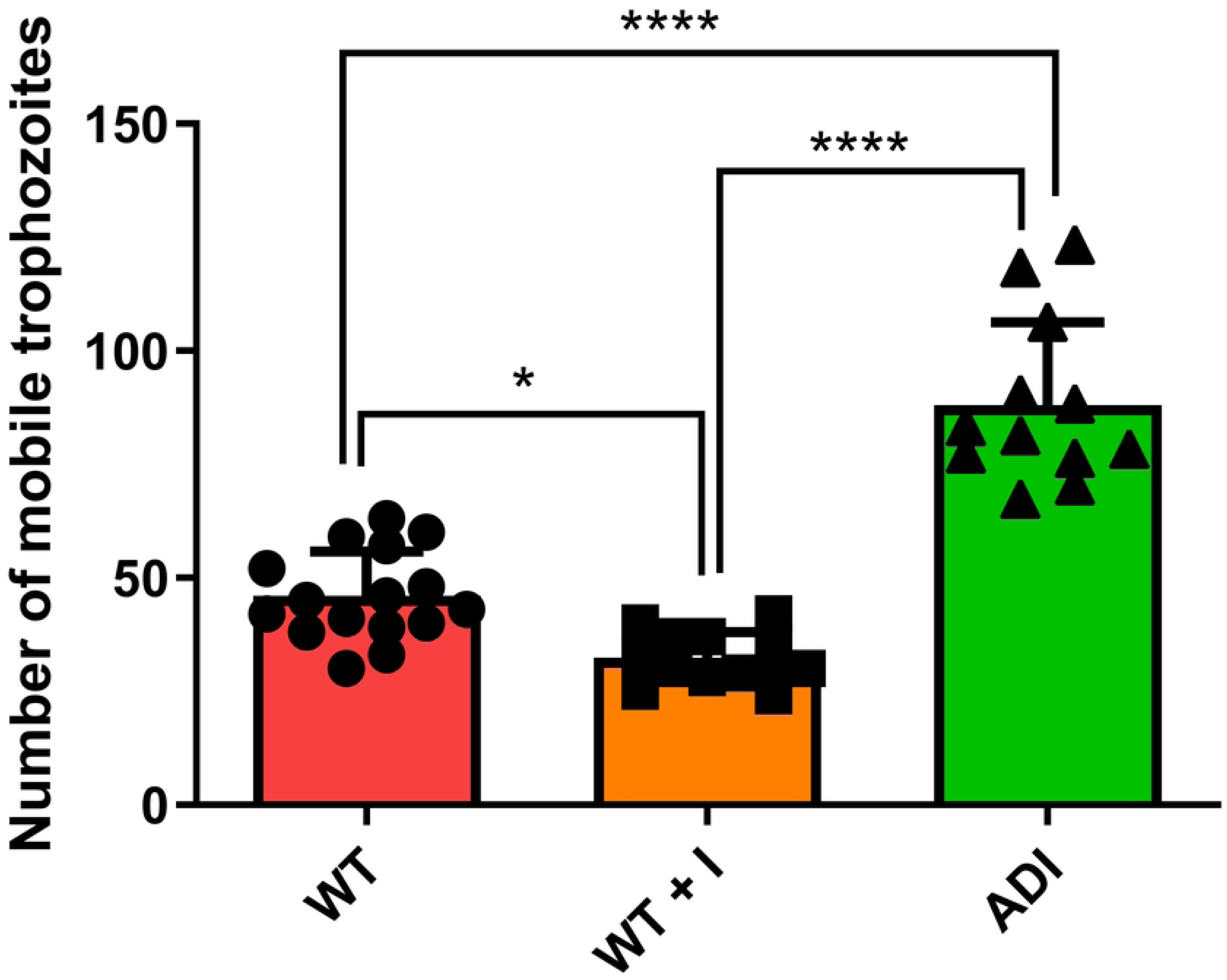
Indole affects F-actin production, size, and mobility in *E. histolytica* trophozoites. (A) Confocal microscopy images (X20) of *E. histolytica* cells. Two-dimensional images showing bright-field, stained with phalloidin (in red), showing F-actin, merged image, and three-dimensional confocal microscope images of WT, TI, and ADI cells stained with Phalloidin. The scale bar represents 30 µm. (B) F-actin intensity from three-dimensional images using Imaris software. (C) Cell volume analyzed from three-dimensional images of WT, TI, and ADI based on confocal microscopy images. (D) Number of motile trophozoites (WT, TI, and ADI) that went through the 8 µm pore membrane in Transwell assays. Statistical analysis was performed using a one-way ANOVA test, with significance levels denoted as * for a *p*-value <0.05, and **** for a *p*-value <0.0001. The data represent the average results obtained from two independent biological replicates.

To directly assess trophozoite motility, a transwell migration assay was conducted. This assay quantifies the ability of trophozoites to migrate through 8 μm pores in a membrane, thereby assessing their motility under different conditions (WT, WT + I, and ADI). The mobility in ADI increased by 42% compared to WT (Fig 6D). Although WT + I cells exhibited an equivalent cell volume to WT cells (Fig 6B), fewer WT + I cells migrated to the lower chamber compared to WT (Fig 6D).

### Effects of indole on virulence markers and pathogenic behavior

Proteomics data indicate that indole upregulates the level of known virulence factors and proteins impacting *E. histolytica* virulence, like some EhCPs [46], Gal/Gal lectin [48], protein kinases [47], or eukaryotic translation initiation factor 5A [43]. To evaluate the impact of indole on *E. histolytica* virulence, we first performed in vitro cytopathic and bacterial biofilm degradation assays. The results showed that the ADI strain exhibited a 2-fold increase in both cytopathic activity and a 1.5-fold increase in biofilm degradation compared to the WT and WT + I strains (Fig 7A and B). Next, the effect of indole in the mouse model of amebic colitis was investigated. *E. histolytica* establishes a persistent intestinal infection when trophozoites are injected into the mice’s cecum [9].

**Figure 7:**
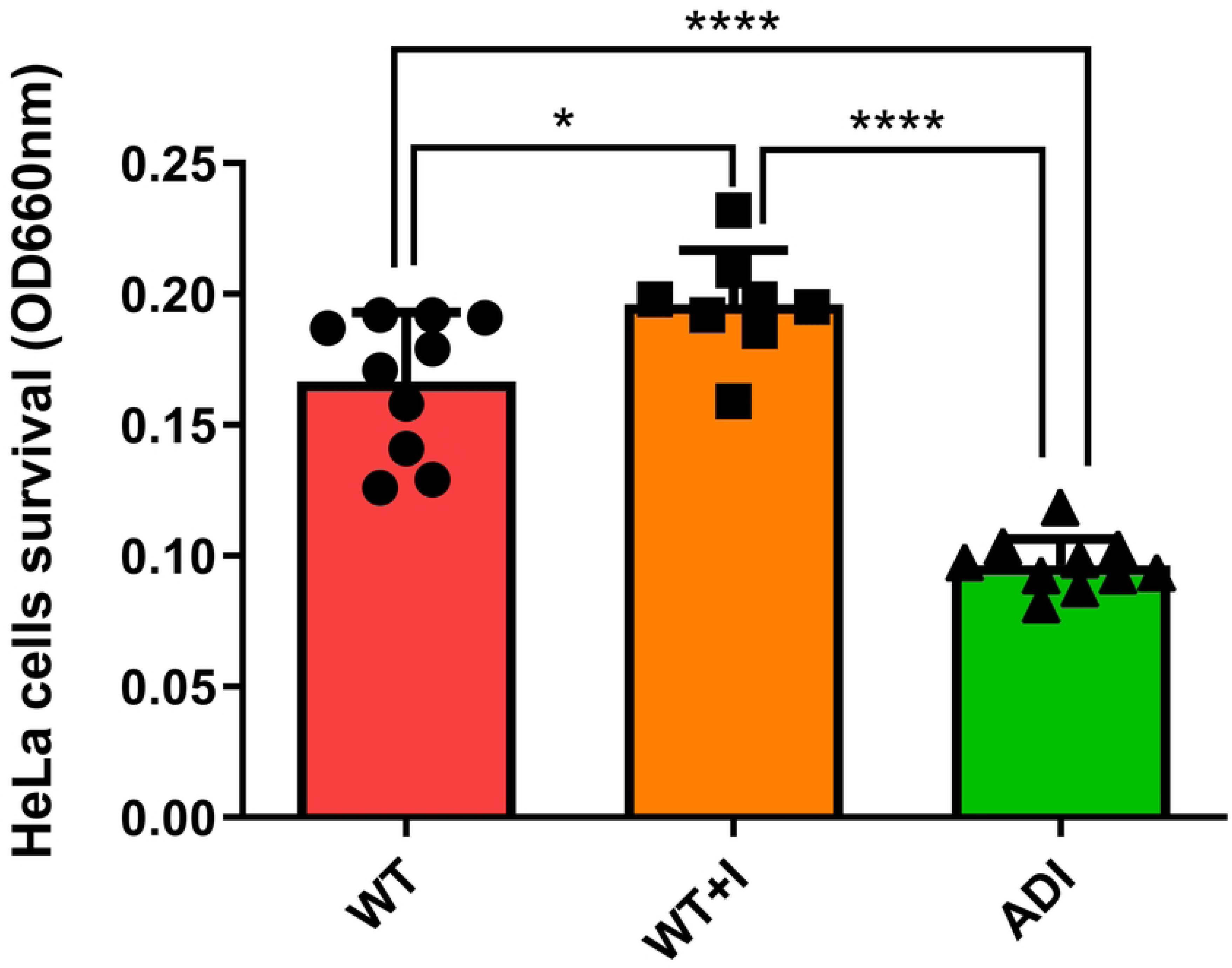
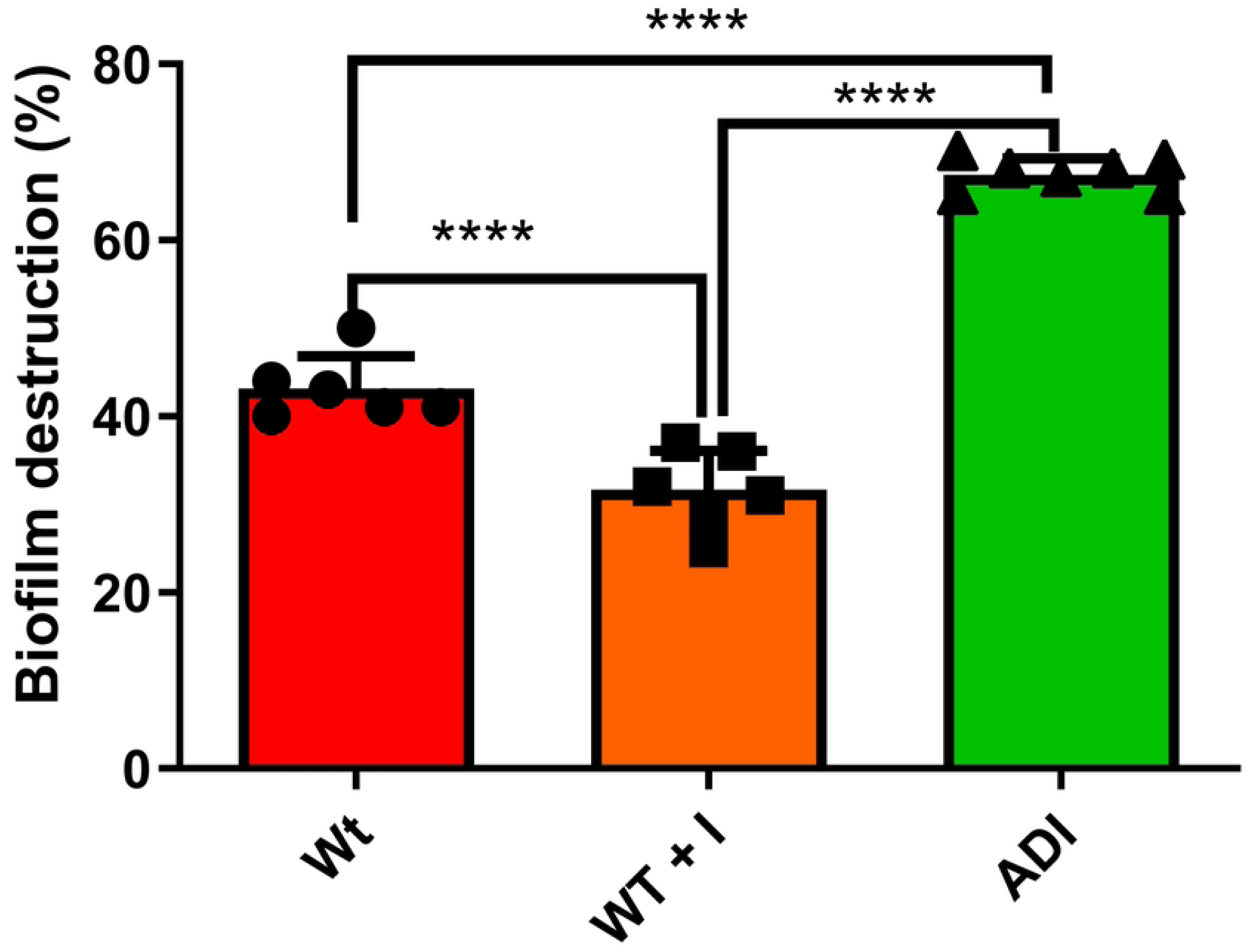
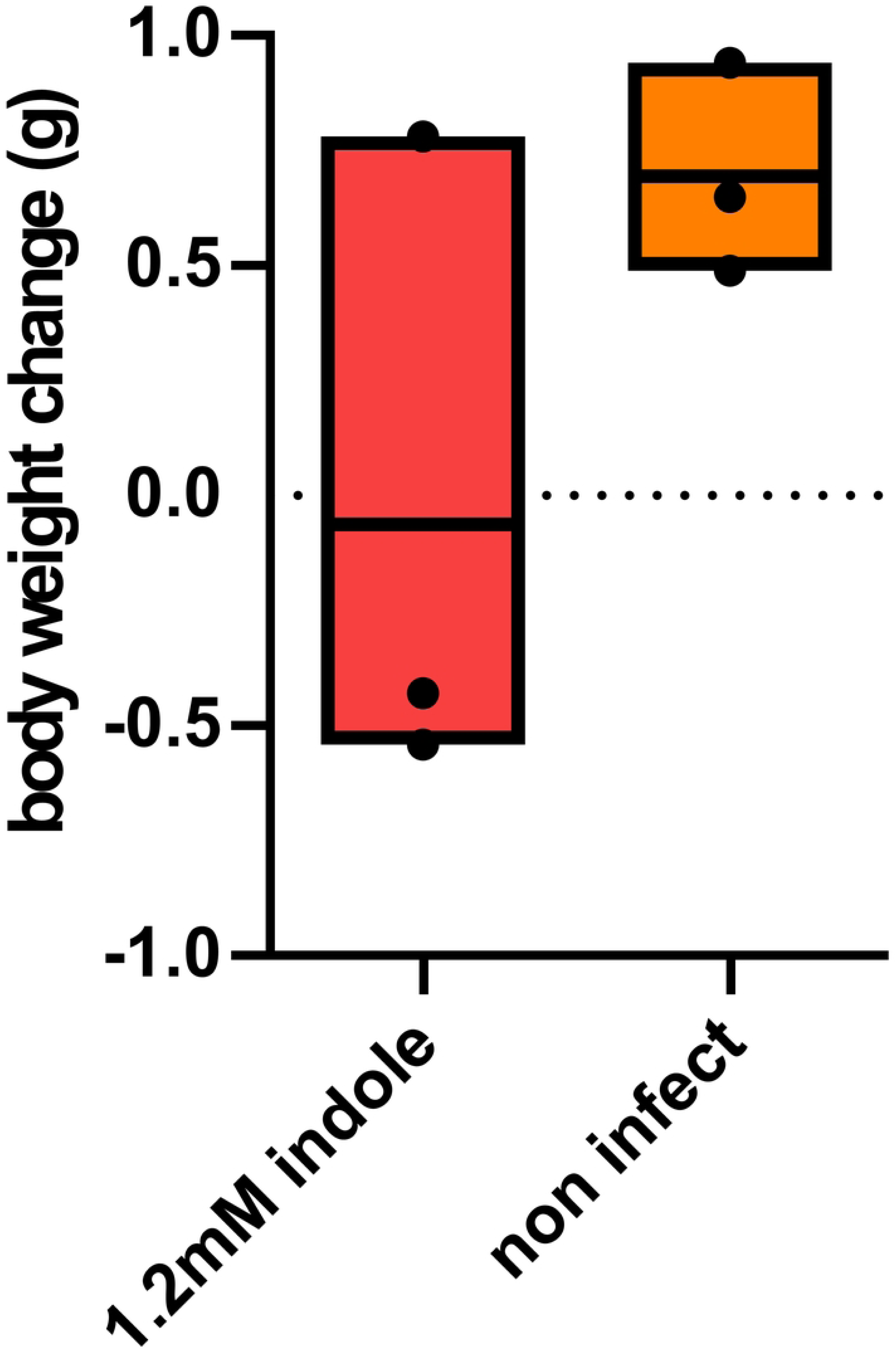
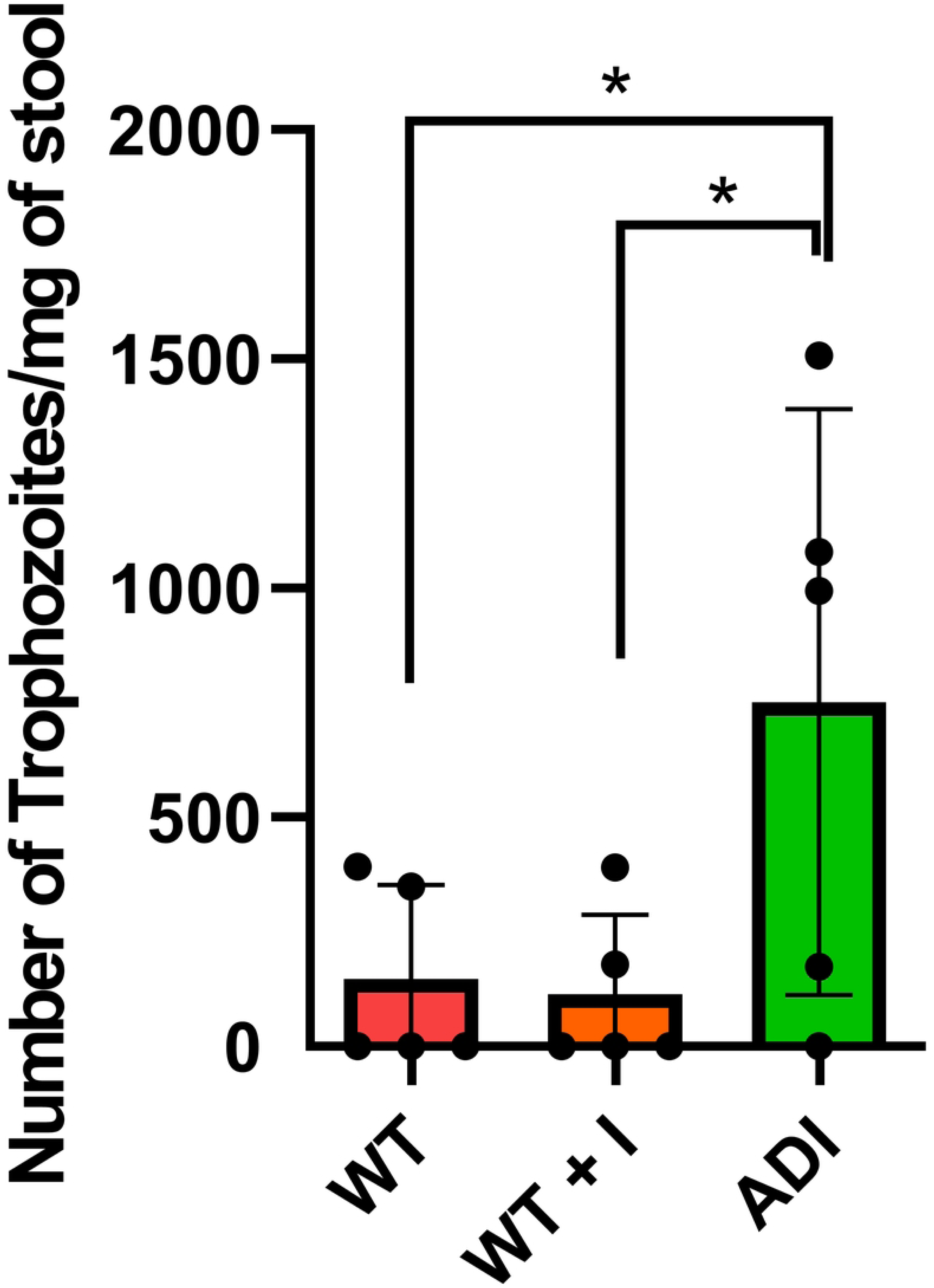
Indole modulates virulence-associated phenotypes of *E. histolytica* in vitro and in vivo. (A) Trophozoites (3*10^5^ cells) under different conditions (WT, WT+I, ADI) were incubated with HeLa cells for 30 minutes. Surviving HeLa cells still attached to the plate were quantified with methylene blue staining. (B) *B. subtilis* biofilm degradation after 3h incubation at 37 °C by 10^6^ trophozoites under different conditions (WT, WT+I, and ADI). (C) Mouse weight control, or with the injection of 1.2 mM indole after 7 days. (D) Number of trophozoites per mouse stool seven days after trophozoite injection in the mouse cecum. Statistical analysis was performed using a one-way ANOVA test, with significance levels denoted as **** for a *p*-value <0.0001. The data represent the average results obtained from two independent biological replicates.

Trophozoites of each strain (WT, WT + I, and ADI) were injected into the mouse cecum, and after seven days of post-infection, the colonization of the parasite in the mouse cecum was determined by quantifying the amount of *E. histolytica* 18S rRNA in the stool. We first evaluated the effect of indole (1.2 mM) on body weight in mice and found no significant changes (Fig 7C). Mice were then injected with WT, WT + I, and ADI trophozoites. Seven days post-inoculation, the number of trophozoites in the stool was determined to assess parasite colonization. The number of ADI trophozoites per mg of stool was 2-fold higher than that of both WT and WT + I (p < 0.05, Fisher’s LSD) (Fig 7D). Unlike ADI trophozoites, which significantly colonize the mouse cecum compared to the WT, WT + I trophozoites showed no such increase.

## Discussion

The presence of indole, a bacterial metabolite, in the gut has emerged as a significant modulator of *E. histolytica* biology. Our study demonstrates for the first time that *E. histolytica* not only internalizes exogenous indole but also adapts to its cytotoxicity. We observed a clear accumulation of indole within the parasite when it was provided extracellularly, indicating that indole enters the cell. It is now widely accepted that indole, in both prokaryotic and eukaryotic systems, can diffuse passively across biological membranes without the need for a specific protein transporter [40, 48]. Interestingly, the enzyme responsible for indole production, tryptophanase, is less abundant in ADI compared to WT trophozoites, suggesting that *E. histolytica* reduces its endogenous indole synthesis when indole is already present in the environment. To our knowledge, tryptophanase repression by indole itself has not been documented. However, in bacteria, tryptophanase is known to be subject to catabolite repression in the presence of glucose, a more favorable carbon source [49]. This suggests that *E. histolytica* may have evolved a distinct regulatory response to intracellular indole accumulation.

This adaptation results in a tolerance for elevated intracellular indole concentrations and leads to a stress-resistant and more virulent phenotype. These findings suggest that indole, beyond acting as a stressor, may function as a microbial signal that primes the parasite for survival and enhances its pathogenic potential within the host. The gut represents a natural environment where *E. histolytica* is likely to encounter indole, with concentrations ranging from 0.3 to 6.6 mM in humans and 200 to 400 μM in the mouse cecum [19] [50]. Therefore, the trophozoite’s adaptation to indole may be critical for effective colonization and virulence *in vivo*.

Proteomic analysis reveals two major responses of *E. histolytica* to indole adaptation. The first is increased levels of actin-related proteins associated with enhanced motility and virulence, and the second is the elevated resistance to oxidative stress (OS). These adaptation mechanisms lead to more virulent ADI trophozoites.

The enhanced motility observed in ADI trophozoites correlates with the upregulation of actin protein, EHI_043640, α-Actinin-2, EHI_161200, and regulators of actin dynamics, such as RhoGAPs and RhoGEFs. Actin protein forms dynamic filaments that shape the cell and enable movement. α-Actinin-2 stabilizes the cytoskeleton by crosslinking actin filaments [51]. RhoGEFs activate Rho GTPases, promoting actin polymerization and structure formation, while RhoGAPs inactivate them, leading to actin disassembly. Together, these proteins tightly regulate cytoskeletal dynamics [52]. Notably, the actin protein EHI_043640 is a truncated version of EHI_182900 with an identical amino acid sequence. Although its precise functional role remains undetermined, its selective expression under indole exposure suggests it may participate in adaptive cytoskeletal regulation. It is highly upregulated in virulent *E. histolytica* following human colon invasion [53] and in ADI trophozoites.

Beyond molecular adaptations, ADI trophozoites exhibit a smaller cell size, which facilitates amoeboid movement. In this migration mode, characterized by limited frictional drag and low traction forces, a reduced cell body size helps minimize resistance and enhances motility [54]. This physical advantage likely enables trophozoites to navigate more efficiently through dense and heterogeneous environments, such as the intestinal mucus layer or host tissues. A similar phenomenon has been observed in normal human breast cells, which, upon transformation into tumor cells following exposure to transforming growth factor β, exhibit reduced size, increased migration speed, and decreased contractile forces [55].

Importantly, adaptation to indole leads to a more virulent phenotype, as evidenced by increased cytotoxicity toward HeLa cells, enhanced biofilm degradation and more efficient colonization of the mouse cecum. This is somewhat paradoxical, given that certain classical virulence factors, such as EhCPs [56] and β-amylases [57], are downregulated in ADI. Specifically, CP1, CP4 (EHI_050570), and CP6 (EHI_151440) are reduced, while others like CP7 (EHI_039610) and EHI_010850 are upregulated. Interestingly, CP4 and CP6 are typically more abundant in the virulent HM-1:IMSS strain, whereas CP7 is enriched in the less virulent Rahman strain [58]. This suggests that CP4 and CP6 may not be essential for virulence in the context of indole adaptation, making ADI a unique example where virulence is maintained or even enhanced despite downregulation of canonical CPs.

Similarly, β-amylases [51], which are important for mucus degradation and host colonization, are downregulated in ADI. Their expression is known to be suppressed in the presence of bacteria [58], suggesting that indole may act as a microbial signal indicating bacterial abundance and reduce the need for amylase-dependent digestion. In contrast, other virulence-associated proteins, such as Gal/GalNAc lectin [59] and actin [60] are upregulated in ADI, offering alternative pathways for tissue invasion and host interaction. Together, these findings suggest that changes in cytoskeletal dynamics and physical properties, particularly enhanced motility and reduced drag, may compensate for the downregulation of some classical virulence factors. We propose that the actin-driven motility enhancements observed in ADI trophozoites play a central role in their increased cytopathic activity, biofilm degradation, and successful colonization of the mouse cecum. This idea is supported by earlier studies showing that pharmacological disruption of the actin cytoskeleton in *E. histolytica*, through agents such as cytochalasin D, leads to impaired motility, reduced host cell killing, biofilm degradation (Ankri, *unpublished*) and attenuated virulence in vivo [61, 62].

Infection with *E. histolytica* has been shown to increase the abundance of *Prevotella copri*, which in turn depletes the endogenous indole derivative indole-3-pyruvate (IPyA). Since IPyA induces T regulatory cell 1 (Tr1) and strengthens the intestinal immune barrier, its reduction exacerbates intestinal inflammation and the symptoms of amebic diarrhea [67][68][69]. Although our current experiments did not measure inflammation levels in infected mice, this interplay between *E. histolytica*, microbiota-derived indole metabolites, and host immunity represents an important avenue for future investigation.

Adaptation to indole in *E. histolytica* leads to upregulation of translational machinery components, suggesting increased protein synthesis. However, the SUnSET assay showed no significant difference in overall protein synthesis between ADI, WT, and WT+I strains. This discrepancy may be explained by two factors: first, proteomics provides a snapshot of protein levels in the parasite, reflecting the balance between synthesis and degradation, whereas SUnSET specifically measures newly synthesized proteins. Second, proteomics identified upregulation of specific protein subsets (94 and 100 proteins increased in ADI versus WT and WT+I), while SUnSET assesses total protein synthesis and may not be sensitive enough to detect changes limited to particular proteins.

Resistance to OS is a prominent feature of ADI trophozoites. Proteomic analysis reveals upregulation of oxidoreductases and other OS-response proteins, and functional assays confirm that ADI trophozoites survive better upon H₂O₂ exposure. This enhanced OS resistance aligns with prior findings linking OS resilience to increased virulence in *E. histolytica* [63]. For example, the virulent HM-1:IMSS strain expresses higher levels of OS-response genes and shows greater survival under H₂O₂ challenge than the avirulent Rahman strain [64, 65]. Interestingly, ADI and control trophozoites exhibit similar levels of oxidized proteins following H₂O₂ exposure. This apparent paradox may be explained by efficient proteostasis mechanisms in ADI trophozoites. Upregulation of the protein translation machinery likely reflects an increased demand for new protein synthesis, especially as a compensatory response to oxidative damage. ADI cells show elevated levels of heat shock protein 70 (Hsp70, EHI_150770), a chaperone involved in proteostasis and degradation of oxidized proteins. Thus, although the absolute amount of oxidized proteins remains comparable among WT, WT + I, and ADI, accelerated turnover driven by Hsp70 and related factors may maintain protein homeostasis under oxidative stress [66].

While the molecular mechanisms behind indole adaptation are likely multilayered and complex, our study (summarized in Fig 8) reveals a coordinated cellular strategy deployed by *E. histolytica* to face this bacterial metabolite. This involves remodeling the cytoskeleton and enhancing oxidative stress resistance to meet new physiological demands.

**Figure 8:**
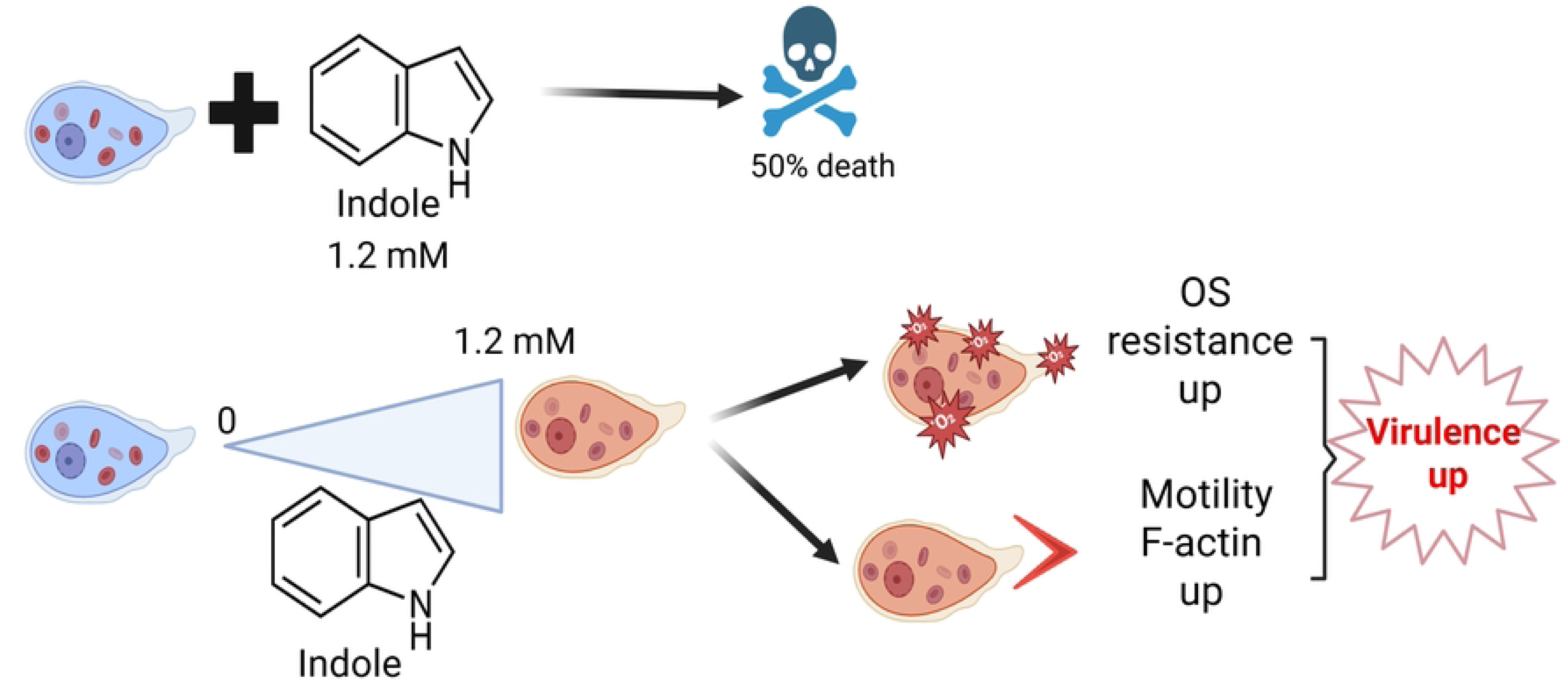
Summary: Indole exposure kills *E. histolytica*, but gradual adaptation leads to enhanced OS resistance, motility, and virulence. The Figure was created in BioRender. Zanditenas, E. (2025) https://BioRender.com/p7y4caz.

In conclusion, this work uncovers how a common gut metabolite can reshape the physiology and virulence of a human parasite. It highlights the remarkable plasticity of *E. histolytica*, which not only tolerates but thrives in the presence of microbial signals such as indole. These findings open up new perspectives on how microbiota-derived metabolites might influence parasite behavior and disease outcome, an emerging field that clearly deserves further exploration.

## Acknowledgments

We appreciate the support of the Laura and Isaac Perlmutter Metabolomics Center, a part of the Biomedical Core Facility. We would like to thank Bella Agranovich and Ifat Abramovich for their assistance in metabolomics experimental design, sample processing, and data analysis.

We appreciate the help of Maya Holdengreber with the software Imaris for microscopy images analysis and the help of Ziv Tamar and Renana Fischer-Weinberger for proteomics data analysis. We thank Prof. Michael Lalk (Greifswald University, Germany) for his critical reading of this manuscript. We also acknowledge the Alfried Krupp von Bohlen und Halbach Foundation (Greifswald, Germany) for providing an optimal environment to S.A. during the preparation of this manuscript.

---

**S1 Appendix: Indole affects F-actin production and size of *E. histolytica* trophozoites**. Quantification of the (A) F-actin intensity fromt-dimensional images, using Cellpose-SAM model coupled with Fiji. (B) Quantification of cell area for the two-dimensional images. Statistical analysis was performed using a one-way ANOVA test, with significance levels denoted as * for a *p*-value <0.05, and **** for a *p*-value <0.0001. The data represent the average results obtained from two independent biological replicates.

**S1 Table:** Proteomics data showing all proteins up and down regulated in trophozoites (WT, WT+I, and ADI) detected by mass spectroscopy. The proteins were cleaved with trypsin and analyzed by the LC/MSMS using the Exploris480 (Thermo) mass spectrometer in DIA mode in order to increase the number of identifications. The data was analyzed using DIA-NN 1.9.1 for identification and quantification against Entamoeba histolytica proteome from Uniport with 1% FDR threshold (FDR =false discovery rate, is the estimated fraction of false positives in a list of peptides). Additional statistical analysis was done by Perseus 1.6.7.0. The protein list contains the quantitation values, peptide count, T-test and annotations columns. The significantly changed proteins are marked as detailed in the color legend on the top rows. Difference is log 2 (fold changed), Log2(0) in log2 intensity columns was replaced with 13 (the threshold intensity in the project). A protein identified with a single peptide could not be considered as a certain identification marked in red in column called “# of peptides” (column AF).

